# A Pluripotent Stem Cell Platform for in Vitro Systems Genetics Studies of Mouse Development

**DOI:** 10.1101/2024.06.06.597758

**Authors:** Rachel A. Glenn, Stephanie C. Do, Karthik Guruvayurappan, Emily K. Corrigan, Laura Santini, Daniel Medina-Cano, Sarah Singer, Hyein Cho, Jing Liu, Karl Broman, Anne Czechanski, Laura Reinholdt, Richard Koche, Yasuhide Furuta, Meik Kunz, Thomas Vierbuchen

## Abstract

The directed differentiation of pluripotent stem cells (PSCs) from panels of genetically diverse individuals is emerging as a powerful experimental system for characterizing the impact of natural genetic variation on developing cell types and tissues. Here, we establish new PSC lines and experimental approaches for modeling embryonic development in a genetically diverse, outbred mouse stock (Diversity Outbred mice). We show that a range of inbred and outbred PSC lines can be stably maintained in the primed pluripotent state (epiblast stem cells -- EpiSCs) and establish the contribution of genetic variation to phenotypic differences in gene regulation and directed differentiation. Using pooled *in vitro* fertilization, we generate and characterize a genetic reference panel of Diversity Outbred PSCs (n = 230). Finally, we demonstrate the feasibility of pooled culture of Diversity Outbred EpiSCs as “cell villages”, which can facilitate the differentiation of large numbers of EpiSC lines for forward genetic screens. These data can complement and inform similar efforts within the stem cell biology and human genetics communities to model the impact of natural genetic variation on phenotypic variation and disease-risk.

## INTRODUCTION

Large-scale genome-wide association studies (GWAS) in human populations have now identified variants at tens of thousands of genomic loci that modulate the risk of developing a variety of common diseases^1,2^. Recent evidence also suggests that a significant subset of GWAS variants contribute to adult phenotypic variation by altering developmental processes^3–8^. However, most experimental model systems for studying embryonic development, such as inbred mouse strains, tend to have very limited genetic diversity, and as a result, we have a relatively limited understanding of how natural genetic variation contributes to phenotypic variation in mammalian development. The lack of tractable experimental models for studying embryonic development in genetically diverse populations thus represents an important gap in the field.

There is an increasing appreciation that studies performed in genetically diverse, outbred individuals have an increased likelihood of generating results that generalize to other biological contexts^9–13^. For example, it is well established that the penetrance of phenotypes observed in genetically engineered mouse models (e.g. gene knockouts and models of human disease) can be highly dependent on the genetic background used for the study^14–16^. In addition, genetically diverse populations can be used for forward genetic approaches to understand the genetic architecture of complex traits and map sequence variants (quantitative trait loci, QTL) contributing to phenotypic variation at the molecular, cellular, and organism scale^17–20^. Thus, experimental systems that incorporate genetic variation will be essential for characterizing genotype-phenotype relationships in complex biological systems and for generating more robust pre-clinical models of human disease biology.

These observations have stimulated intense interest in characterizing the impact of genetic variation on phenotypic variation between individuals during embryonic development^21–24^. For example, mapping QTLs that contribute to functional variation in molecular and cellular traits in developing tissues can complement traditional reverse genetic approaches and genome-wide loss-of-function genetic screens to better elaborate fundamental biological processes^19,25–29^. However, QTL mapping studies of developmental processes *in vivo* have proven difficult to design and scale since for many phenotypes each genetically distinct individual can only be assayed at a single developmental stage, making it difficult to ascertain how genetic variation manifests as functional variation during the dynamic process of tissue development. In recent years, there has been increasing interest in using directed differentiation of pluripotent stem cells (PSCs) *in vitro* as a model for studying natural genetic variation in developing cell types^17,18^. PSC differentiation models can be used to generate a variety of disease-relevant cell types and recapitulate key features of embryonic developmental processes, enabling experimental designs (such as longitudinal studies) that are not practical *in vivo*^30,31^.

In this study, we combine the power of pluripotent stem cell models with the genetically diverse Diversity Outbred (DO) mouse stock. The DO stock is a highly recombinant intercross population generated by crossing five commonly-used inbred mouse strains (129S1/SvImJ, A/J, C57BL/6J, NOD/ShiLtJ, NZO/HiLtJ) and three genetically divergent wild-derived inbred strains (CAST/EiJ, PWK/PhJ, WSB/EiJ)^32^. The DO stock encompasses significant genetic and phenotypic heterogeneity, capturing ∼90% of the genetic variation present across common inbred mouse strains. The DO population contains over 40 million single nucleotide polymorphisms (SNPs), has a high frequency of heterozygosity across the genome, but maintains high minor allele frequencies (>5%) for most SNPs (**Fig. 1A**)^33,34^. These features have enabled high-resolution QTL mapping studies for a number of complex phenotypes including animal behavior, physiology, aging, and cancer metastasis^35–40^. More recently, efforts have been made to establish and characterize PSC panels from DO founder strains and outbred DO mice, and to map QTLs in naïve pluripotent stem cells, opening up the possibility of extending these approaches to developing cell populations differentiated from PSCs^25,41–43^. However, current resources and approaches for performing large-scale *in vitro* QTL studies with DO PSCs are limited.

**Figure 1.**
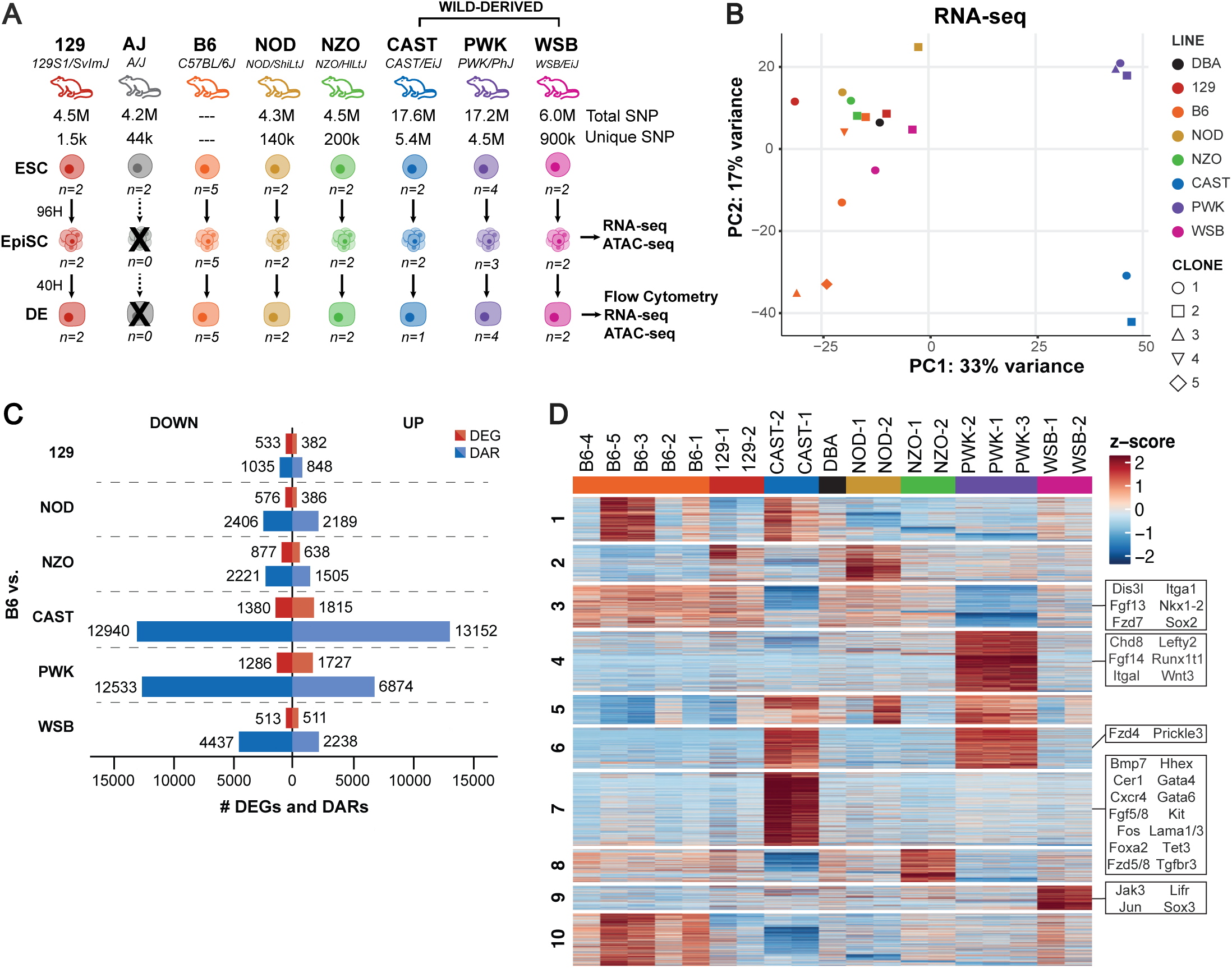
Strain survey of DO founder EpiSCs reveals contribution of genetic diversity to transcriptional and epigenomic heterogeneity in primed pluripotency. **(A)** Overview of strain survey experimental design. Total and unique single nucleotide polymorphisms (SNPs) of each strain compared to C57BL/6 (reference genome). A minimum of two, independently derived ESC lines from each of the 8 DO founder strains were converted to EpiSCs and characterized by bulk RNA- and ATAC-sequencing. Directed differentiation of EpiSCs into definitive endoderm was also assessed (differentiation efficiency, RNA-seq and ATAC-seq of MACS-purified DE). **(B)** PCA of gene expression (RNA-seq) in EpiSCs from each DO founder strain. DBA/2J EpiSCs are included for comparison. **(C)** Total number of differentially expressed genes (DEGs) and differentially accessible regions (DARs) in DO founder EpiSCs compared to B6 EpiSCs (FDR ≤ 0.1, absolute fold change > 2). **(D)** k-means clustering (k =10) of RNA-seq data from DO founder EpiSCs. For selected clusters, differentially expressed genes related to transcriptional regulation and developmental signaling pathways are highlighted.

Here, we develop novel experimental approaches and cellular resources to perform population-scale *in vitro* directed differentiation of PSCs from DO mice. To establish the contribution of genetic variation to phenotypic variability in early development, we characterize transcriptomic, epigenomic, and functional differences in epiblast stem cells (EpiSCs) and early germ layer progenitors (definitive endoderm – DE) from the DO founder strains. This strain survey suggests a strong genetic contribution to phenotypic variability at these stages, indicating that genetic variants underpinning this phenotypic variation can be mapped in the DO outbred stock. Next, we implemented pooled *in vitro* fertilization (IVF) to derive genetically diverse DO PSC lines at scale. We find that genetically distinct cell lines from both male and female DO mice (n=230) can be stably cultured in the primed pluripotent state, overcoming a significant hurdle for the generation of large cell panels for QTL mapping. To enable population-scale experiments, we demonstrate the feasibility of multiplexed cell culture approaches (“cell villages”) and implement a computational method (Dropulations) for genetic demultiplexing of single-cell RNA-sequencing (scRNA-seq) to characterize DO EpiSC villages. This new platform opens up the possibility of *in vitro* population genetics studies to dissect fundamental mechanisms of cellular differentiation, extending upon what is possible with analogous studies using human PSCs^27,28,44,45^.

## DESIGN

Our goal was to develop a PSC platform from DO mice that can facilitate QTL mapping in the context of *in vitro* differentiation models. Our rationale for choosing to use PSCs from the DO stock, rather than currently available hPSC panels from different individuals, was based on several considerations: 1) The DO stock is a recombinant intercross population that was generated specifically for mapping the genetic basis of complex traits, such that far fewer individuals are needed to perform sufficiently powered QTL mapping studies relative to human PSC panels, 2) hPSC panels generally provide a very limited representation of human genetic diversity (i.e. they currently consist predominantly of individuals of European descent), are costly to acquire, and subject to more restrictive regulation due to ethical considerations, making it difficult for most labs to routinely work with these panels, 3) Results from mouse PSC models can be rigorously evaluated *in vivo*, whereas direct validation of findings from human PSC models remain difficult or impossible in many cases.

However, there were important hurdles to overcome to establish PSC panels from the Diversity Outbred stock as an experimental platform for performing QTL mapping studies. For example, compared to hPSCs, the culture and directed differentiation protocols available for genetically diverse mouse PSCs have not been as extensively optimized. To establish a DO platform that can be used for QTL mapping studies, several limitations needed to be addressed, including (1) identifying culture conditions that support consistent culture and directed differentiation of genetically diverse mouse PSCs, (2) deriving large enough panels of DO PSC lines for sufficiently powered QTL mapping studies (>200 PSC lines), and (3) developing approaches that can facilitate consistent *in vitro* differentiation of large numbers of DO PSC lines. We and others have demonstrated that mouse EpiSCs can be cultured stably in media containing Wnt pathway inhibitors and are a better starting point for directed differentiation studies compared to naïve pluripotent stem cells^46–51^. We reasoned that culturing DO PSCs in the primed pluripotent state (EpiSCs) might enable more routine culture and differentiation of genetically diverse PSC lines.

## RESULTS

### Generation of epiblast stem cell lines from DO founder strains

We first sought to determine whether PSCs from each of the eight Diversity Outbred founder strains can be stably cultured in the primed pluripotent state (EpiSCs) (**Fig. 1A**). We acquired a well-characterized panel of ESC lines from each of the eight Diversity Outbred founder strains (n = 2 independently-derived ESC lines / inbred strain) and performed *in vitro* ESC-EpiSC conversion of each DO ESC line (see methods; **Table S1**)^25,52^. We were able to stably convert most (n= 18/20) of the ESC lines into EpiSC lines that can be maintained on irradiated feeders in media containing FGF2, Activin A, and a small molecule inhibitor of the canonical Wnt signaling pathway (NVP-TNKS656) (see methods; **Fig. S1A**). Surprisingly, both ESC lines from the A/J strain exhibited significant cell death and poor cell attachment during EpiSC conversion and thus could not be converted into stable EpiSC lines, despite multiple attempts (**Fig. S1B**). Similar results were obtained from an independent set of clonal A/J iPSC lines (n = 24 subclones), indicating that the genetic background is likely responsible for this issue rather than a specific technical problem with the original two A/J ESC lines (**Fig. S1C-D**). Simple modifications to the EpiSC culture protocol (supplementation with Vitamin C, passaging the cells as larger colonies) slightly improved the growth of A/J EpiSCs, but were not sufficient to stably culture these lines beyond three passages (**Fig. S1C-D**).

These results indicate that primed pluripotency culture conditions support the stable culture of EpiSCs from 7/8 DO founder strains and provide an initial indication that genetic variation present across the DO founder strains can contribute to significant phenotypic variation in EpiSCs.

### Strain survey of gene expression and chromatin accessibility in EpiSCs from DO founder strains

Having established EpiSC lines from seven of the DO founder strains, we next sought to characterize variation in gene expression (bulk RNA-sequencing) and cis-regulatory element activity (assay for transposase-accessible chromatin with high-throughput sequencing [ATAC-seq]) among these lines (**Fig. 1A**). Initial inspection of gene expression data confirmed that EpiSCs from each strain exhibit transcriptional signatures of primed pluripotent stem cells, with high expression of general (*Pou5f1*, *Sox2*, *Nanog*) and primed (*Otx2*, *Pou3f1*) pluripotency regulators and low expression of naïve pluripotency markers (*Klf2, Nr5a2, Tbx3)* and genes associated with germ layer differentiation *(Sox1, Sox17, Kdr, Gata6)* (**Fig. S2A**). We did observe some variation in gene expression between strains, particularly among the CAST/EiJ lines which have increased expression of genes associated with spontaneous differentiation (*T, Cer1, Cdx2*) (**Fig. S2A**).

Unsupervised principal component analysis (PCA) of gene expression (RNA-seq) and chromatin accessibility (ATAC-seq) data revealed that EpiSC lines clustered predominantly by their strain of origin, indicating that genetic differences between strains are a major driver of molecular variation between lines (**Figs. 1B, S2B**). There was also a general correlation between genetic divergence (the number of SNPs between lines) and transcriptional/epigenetic variation, with the wild-derived inbred strains CAST/EiJ and PWK/PhJ clearly separated from the more closely related common laboratory strains along PC1/PC2 in both RNA-seq and ATAC-seq PCA plots (**Figs. 1B, S2B**). Accordingly, CAST/EiJ and PWK/PhJ EpiSCs also exhibited a higher number of differentially expressed genes (FDR≤0.1, absFC≥2) and differentially accessible cis-regulatory elements (FDR≤0.1, absFC≥2) compared to C57BL/6J than were observed in pairwise comparisons between C57BL/6J EpiSCs and the other common laboratory strains (**Fig. 1C**).

To further investigate the specific transcriptional and epigenomic features that distinguish each of the strains, we performed k-means clustering on both the RNA-seq (**Fig. 1D**) and ATAC-seq (**Fig. S2C**) datasets. This analysis identified clusters of genes (e.g., cluster 3 and 6) whose expression differs between the wild-derived inbred strains CAST/EiJ and PWK/PhJ and the common laboratory strains, as well as genes that are consistently differentially expressed across each of the individual strains. Examination of the specific genes that distinguish EpiSCs from each strain revealed increased expression of genes associated with primitive streak and endodermal differentiation (*Cer1*, *Cxcr4*, *Foxa2*, *Hhex*, *Gata4*, *Gata6*) in CAST/EiJ EpiSCs, suggesting a higher frequency of spontaneous differentiation in these lines. This highlights one issue with performing comparisons between these lines using bulk RNA-seq and ATAC-seq, which will be significantly influenced by differences in the relative amount of distinct sub-populations of cells within the culture (i.e., the relative frequency of spontaneous differentiation).

Together, these data indicate that genetic variation (strain of origin) contributes more to variation in molecular phenotypes than non-genetic and technical factors (i.e., variation between independently derived lines from the same genetic background). This is consistent with previous data from naïve pluripotent stem cells from each of these genetic backgrounds as well as studies in genetically diverse panels of human PSCs ^25,42^. In addition, this transcriptomic and epigenomic strain survey can serve as a comparative atlas for future large-scale analysis of EpiSCs from the Diversity Outbred stock, including mapping of expression QTLs and chromatin accessibility QTLs.

### Genetic variation modulates directed differentiation of EpiSCs to definitive endoderm

Having characterized transcriptomic and epigenomic differences across the DO founder strains, we next sought to determine the extent to which these molecular differences impact the functional capacity of EpiSCs to differentiate into a specific lineage. To evaluate this systematically, we chose to differentiate EpiSCs from each of the DO founder strains into definitive endoderm (DE) using our rapid, two step protocol that has previously been shown to convert EpiSCs into DE with >90% efficiency^46^. Given that we wanted to observe how different lines behave under the same conditions, we did not perform any specific optimization for each EpiSC line. To account for technical variability, we included independently derived lines from each strain (n ≥ 2 EpiSC lines/strain; 18 total EpiSC lines) and performed technical replicates for each cell line (n ≥ 2 technical replicates/EpiSC line) (**Fig. 2A**; see methods). Differentiation efficiency into DE was calculated for each line by quantifying the fraction of cells co-expressing DE markers SOX17 and CXCR4 using flow cytometry.

**Figure 2.**
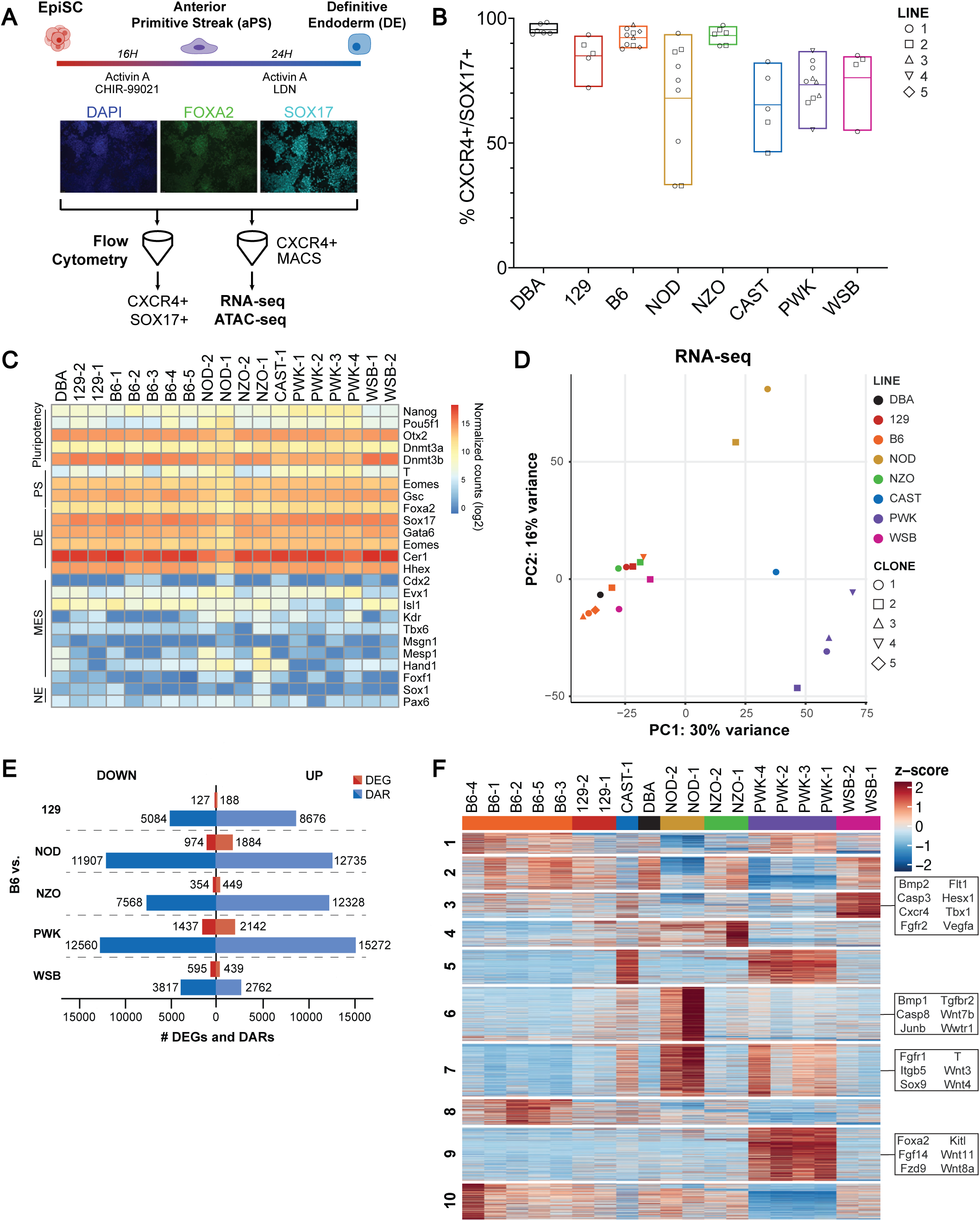
Genetic variation contributes to variation in directed differentiation efficiency and definitive endoderm gene regulation. **(A)** Overview of the definitive endoderm (DE) directed differentiation protocol. A minimum of two EpiSC lines from each of the DO founder strains were differentiated into DE. To determine differentiation efficiency, samples were analyzed by flow cytometry for expression of DE markers CXCR4 and SOX17. To purify DE for bulk RNA-seq and ATAC-seq analysis, CXCR4-positive cells were isolated using magnetic activated cell sorting (MACS). **(B)** Efficiency of directed differentiation into DE (CXCR4+, SOX17+). Directed differentiation efficiency was measured for a minimum of two EpiSC lines from each strain (biological replicates to assess intra-strain variability). In addition, at least two technical replicates (each from different batches of differentiation) were measured from each EpiSC line (indicated by distinct shapes). **(C)** Heatmap of expression of selected marker genes (pluripotency, primitive streak, DE, mesoderm [MES], and neuroectoderm [NE]) in purified DE from each of the DO founder strains. **(D)** PCA of gene expression (RNA-seq) in purified DE from DO founder strains. DBA/2J DE is included for comparison. **(E)** Total number of differentially expressed genes (DEGs) and differentially accessible regions (DARs) in purified DE from each of the DO founder strains relative to B6 (FDR ≤ 0.1, absolute fold change > 2). **(F)** k-means clustering (k=10) of bulk gene expression of purified DE. For selected clusters, differentially expressed genes related to transcriptional regulation and developmental signaling pathways are highlighted.

For each EpiSC line we were able to successfully generate DE, however, there were clear differences in the efficiency of DE specification between different EpiSC lines (range: 98.37% - 33%). Technical replicates of the same EpiSC line and independently derived lines of the same genetic background generally exhibited similar differentiation efficiencies, with the exception of NOD EpiSCs, which gave highly variable results (**Fig. 2B**).

Comparison between strains revealed a consistently lower efficiencies of DE differentiation in EpiSCs from wild-derived strains (CAST, PWK, WSB) compared to the common laboratory strains (129, B6, NZO) (**Fig. 2B**). Taken together, these data indicate that genetic variation contributes to variability in the timing and/or efficiency of lineage specification *in vitro*.

Next, we sought to further characterize transcriptomic and epigenomic variation in DE from each inbred strain. To account for variation in differentiation efficiencies between lines, we isolated putative DE cells (CXCR4+) using magnetic activated cell sorting (MACS) prior to performing bulk RNA- and ATAC-seq (**Fig. 2A**; see methods). Purified DE from each strain exhibited high expression of markers of definitive endoderm genes (*Sox17*, *Foxa2*, *Cer1*), downregulation of core pluripotency genes (*Pou5f1*, *Nanog*), and low expression of genes associated with mesoderm (*Isl1*, *Mesp1*, *Msgn1*, *Tbx6*, *Kdr*, *Hand1*, *Foxf1*) or neuroectoderm (*Sox1*, *Pax6*) differentiation, indicating that purified CXCR4+ cells represent a relatively pure population of DE (**Fig. 2C**). NOD DE exhibited lower expression of core DE genes and higher expression of genes associated with mesodermal lineages (*Kdr*, *Hand1*, *Mesp1*), suggesting a higher frequency of off-target mesodermal differentiation, consistent with the technical variability in DE specification efficiency.

As in the primed pluripotent state, PCA of bulk gene expression and chromatin accessibility clustered lines by genetic background along PC1/PC2, consistent with a dominant effect of genetic background on transcriptional and epigenetic variation in DE (**Figs. 2D, S3A**). Comparing the cell states, there was a far greater number of differentially accessible chromatin peaks in the differentiated DE compared to EpiSCs, suggesting that chromatin accessibility will continue to diverge from pluripotency through differentiation to other lineages (**Figs. 1C, 2E**). Between backgrounds, PWK/PhJ DE exhibited more differentially expressed genes and differentially accessible regions than were observed in comparisons between DE from the standard laboratory strains and C57BL/6J (**Fig. 2E**). Examination of genes and regions of accessible chromatin that vary across genetic backgrounds by k-means clustering highlighted several genes associated with DE specification that were upregulated in specific strains, including *Cxcr4* in WSB/EiJ (cluster 3) and *Foxa2* in PWK/PhJ (cluster 9) (**Figs. 2F, S3B**). Similar to the EpiSC analysis, primitive streak marker *T* was also somewhat upregulated in the CAST/EiJ line as well as the PWK/PhJ lines; however, the most dramatic upregulation of T was found in the two NOD/ShiLtJ lines (**Fig. 2F**). This finding suggests that off-target differentiation to early mesoderm or delayed commitment to DE may contribute to the variability in directed differentiation efficiency.

These data indicate that genetic variation present across the DO founder strains contributes to phenotypic differences in the primed pluripotent state that impact *in vitro* directed differentiation outcomes. We also establish preliminary assays to assess quantitative variation in differentiation phenotypes in genetically heterogenous panels of EpiSCs derived from the DO stock. In addition, these data provide a comprehensive assessment of the impact of genetic variation across distinct inbred mouse strains on gene regulation in two specific early embryonic cell types (epiblast, DE).

### Scalable generation of ESCs from the Diversity Outbred stock via pooled *in vitro* fertilization

Standard methods of ESC derivation are labor-intensive and low throughput, presenting a barrier to the generation of genetically diverse panels of PSCs from the Diversity Outbred stock that are large enough to perform QTL studies of molecular and cellular phenotypes in PSC models of lineage specification^17,19,20^. In addition, ESC lines derived from a single mating pair will all share ∼50% of their genomes, which reduces power for genetic mapping studies due to relatedness^53^. To overcome each of these issues, we reasoned that a multi-parental *in vitro* fertilization (IVF) approach (“pooled IVF”) could be a practical approach to enable scalable generation of genetically diverse (i.e., non-sibling) PSC lines from the Diversity Outbred stock (**Fig. 3A**).

**Figure 3.**
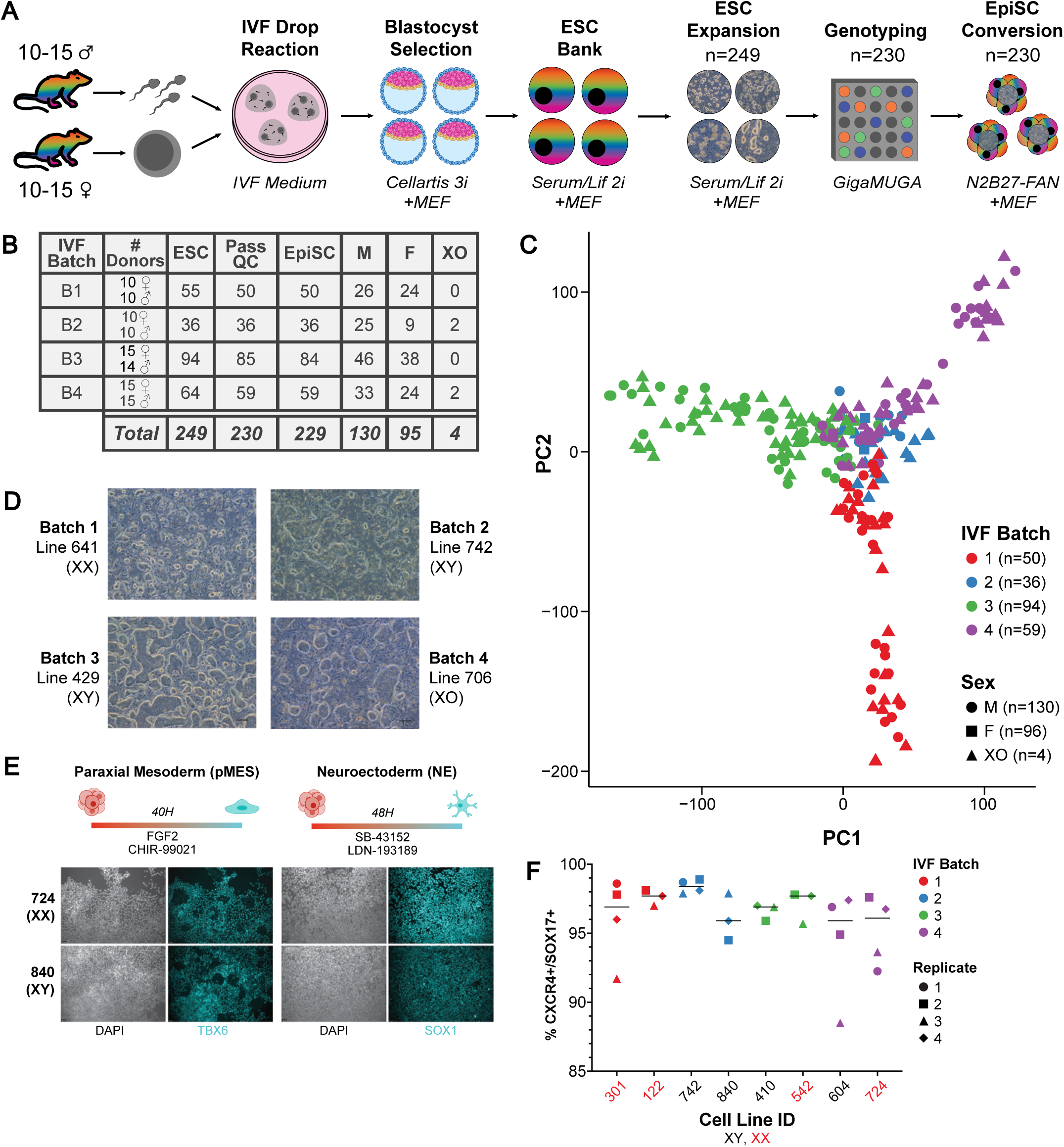
Scalable derivation of DO PSCs using pooled in vitro fertilization. **(A)** Experimental outline of DO ESC derivation using pooled (i.e., multi-donor) *in vitro* fertilization (IVF). **(B)** Summary of DO ESC lines generated using pooled IVF and conversion into EpiSCs. **(C)** PCA of DO PSC genotypes. Samples from distinct batches of IVF are color-coded, and sex genotype is indicated by distinct shapes. **(D)** Representative phase contrast images of early passage DO EpiSCs (p1-2). EpiSC lines derived from ESCs from each of the 4 IVF batches are shown. **(E)** Experimental protocol for directed differentiation experiments (top panel) to assess pluripotency of DO EpiSC lines. Protocols used for paraxial mesoderm (pMES) and neuroectoderm (NE) differentiation are shown. Representative immunostainings (lower panels) for markers of pMES (Tbx6) and NE (Sox1) from individual DO EpiSC lines following directed differentiation. Each differentiation was performed 3 times (3 independent batches of differentiation for each EpiSC line). Qualitatively similar results were obtained from 6 additional DO EpiSC lines. **(F)** Quantification of DE differentiation efficiency from 8 DO EpiSC lines. Definitive endoderm differentiation was performed using the protocol described in Fig. 2A. DE differentiation efficiency (CXCR4+/SOX17+) was quantified using flow cytometry. Each differentiation was performed 3-4 times (3-4 independent batches of differentiation).

To implement this approach, we harvested gametes from non-sibling male and female DO mice (n = 10-15 males, n = 10-15 females for each IVF batch; **Fig 3B**). Sperm and oocytes from all mice were then pooled into several small IVF reactions and successfully fertilized oocytes were allowed to mature into blastocysts. Each blastocyst stage embryo (n = 30-100 embryos/ batch) was then individually isolated for direct derivation of ESC lines *in vitro* or cryo-preserved and then subsequently thawed for ESC derivation (see methods). Based on previous studies deriving ESCs from Diversity Outbred founder strains and DO stock, we chose to use ESC media containing serum, LIF, GSK-3B inhibitor (CHIR99021), and MEK inhibitor (PD0325901) (Serum/LIF/2i) on irradiated fibroblast feeders to facilitate stable maintenance of the naïve pluripotent state^42,52^. Given the potential of prolonged culture of naïve PSCs in media containing 2i to deregulate imprinted genes^54^, we attempted to bank primary stocks of ESCs at low passages (p1-p3). Using this procedure, we derived an initial panel of 249 DO ESC lines from four independent batches of pooled IVF reactions for subsequent experiments (**Fig. 3B**, **Table 1**). In general, most ESC lines from DO mice could be stably maintained in serum/LIF + 2i on feeders and grew well, although there was clearly variability in colony morphology, growth rate, and spontaneous differentiation between DO ESC lines, consistent with observations from previous studies^25,42,43^.

To determine the genotypes of each genetically distinct DO ESC line, we used the Mouse Universal Genotyping Array (GigaMUGA). Importantly, these genome-wide SNP genotyping data also allowed us to determine the sex of each cell line, identify lines with a high-frequency of aneuploid cells, and to identify ESC lines that contain cells from more than one genotype (e.g., from inadvertent transfer of a blastocyst stage embryo that merged/stuck to another embryo prior to isolation for ESC derivation) (see methods). Of the 249 banked ESC lines, 230 ESC lines (130 male, 95 female, 4 ambiguous sex genotype) passed these stringent quality control criteria and were used for downstream experiments.

One potential issue with using a pooled IVF approach is variation in the fertilization efficiency of individual DO males, which could lead to sperm from a small subset of the DO males fertilizing the majority of the eggs in each batch and thus limit the genetic variation captured by the stem cell panel. To assess the diversity of parentage among the DO ESC lines from each batch, we performed kinship analysis of SNP genotype data from DO ESC lines from each of the four batches of IVF. This analysis indicated that most DO ESC lines derived from the same pooled IVF reaction do not share the same parents, confirming that this approach can be used to generate genetically diverse DO PSC panels (**Figs. 3C, S4**). However, there was some variation in the relative amount of ESC lines derived from each sperm/egg donor, with increased relative representation of some individual parents within each batch. Collectively, these data indicate that pooled IVF is a scalable approach for generating genetically diverse stem cell panels from cohorts of Diversity Outbred mice.

### Generation of EpiSCs from Diversity Outbred ESCs

Having generated and characterized a panel of ESCs from DO mice, we next sought to systematically assess whether these ESC lines can be transitioned into the primed pluripotent state *in vitro* and stably maintained as EpiSCs. EpiSCs from the DO stock have not been previously characterized, so it was not clear how they would behave in these culture conditions. We transitioned low-passage, primary DO ESC stocks (n = 230) into EpiSCs using the same approach as what was used for the DO founder strains (see methods; **Fig. 3A**).

Overall, we were able to successfully convert both male (n=130) and female (n=99) lines into stable EpiSC lines in these condition (100% success rate; **Fig. 3B**). Although most EpiSC lines can be stably cultured, we did observe phenotypic variation between EpiSC lines, including differences in colony morphology, spontaneous differentiation (both frequency of differentiation observed and the lineage of cells generated), and re-plating efficiency after splitting (**Fig. 3D)**. These data provide strong evidence that primed pluripotency culture conditions can support the stable growth of cell lines from genetically diverse outbred mice and provide additional evidence (together with data from the DO founder strains; **Fig. 1**) that genetic variation contributes to phenotypic variation in the behavior of primed pluripotent stem cell lines.

Given that EpiSCs from the DO stock have not been previously characterized, we performed additional experiments to assess whether DO EpiSCs are functionally pluripotent, and thus capable of differentiation into each of the three embryonic germ layers. We selected 8 DO EpiSC lines (n=4 males, n = 4 females) and performed directed differentiation into DE, paraxial mesoderm, and neuroectoderm (n = 3 independent batches of differentiation for each cell line; technical replicates). Each of the 8 lines was able to efficiently differentiate into all three embryonic germ layers, confirming that these lines are functionally pluripotent (**Figs. 3E-F**).

Quantification of DE differentiation efficiency demonstrated consistent, high-efficiency differentiation for each of the 8 DO EpiSC lines (**Fig. 3F**). Considered together with the results from definitive endoderm differentiation experiments performed with DO founder strains (**Fig. 2B**), these data demonstrate that this DE directed differentiation protocol works efficiently across a large panel of genetically diverse cell lines (DO founder strain lines + DO EpiSCs; n ∼ 27 total EpiSC lines). In addition, these directed differentiation experiments highlight that by starting from EpiSCs it is possible to achieve rapid and efficient differentiation into progenitors of each embryonic germ layer from a panel of genetically diverse cell lines with no specific optimization performed for each cell line. It would likely be very difficult to achieve similar results starting from ESCs using available protocols.

In summary, we successfully established and banked a panel of genotyped EpiSCs from DO mice (n = 230) that can be shared with the greater scientific community. Taken together, these data provide strong evidence that genetically diverse DO PSC lines from both males and females can be stably maintained in the primed pluripotent state (EpiSCs) and serve as a starting point for rapid and efficient directed differentiation into each of the three embryonic germ layers.

### Generation and characterization of cell villages of DO EpiSCs

Instead of culturing and characterizing each EpiSC line individually, pooled cell culture (“cell villages”) approaches have emerged as a scalable approach for characterizing large panels of genetically distinct cell lines that can reduce technical variation (batch-to-batch variability), significantly lower reagent and labor costs, and enable population-scale perturbation experiments^28,55,56^. For example, this approach has now been used in several studies to analyze cellular phenotypes in genetically diverse human PSC panels^27,45,57^. This approach relies on the fact that each genetically distinct cell line is effectively “barcoded” by its unique genome sequence, allowing for assignment of cells assayed in single cell genomics experiments to specific individuals (genetic demultiplexing)^58^. Having generated a genotyped panel of DO EpiSCs, we next sought to evaluate the possibility of combining sets of these lines into cell villages containing multiple DO EpiSC lines. This could enable large-scale characterization of our DO EpiSC panel (i.e., for quality control purposes) as well as evaluation of molecular phenotypes across DO EpiSCs for quantitative trait locus mapping. However, it has been observed that a few cell lines may rapidly proliferate and overtake a pooled culture due to somatic mutations in tumor suppressors/oncogenes, epigenetic aberrations, aneuploidy, or via active mechanisms such as cell competition^55,59,60^ thereby reducing the utility of this system to screen large numbers of donors, although this has not been observed in all studies employing pooled culture^56,57^. Thus, we also needed to evaluate how well the representation of input cell lines in villages is maintained over time in culture.

To evaluate a cell pooling strategy for DO EpiSCs, we selected 76 DO EpiSC lines (n = 49 male, n = 24 female, n = 3 ambiguous sex genotype) (**Fig. 4A**). To maximize the genetic diversity represented within each cell village (V), we selected DO EpiSCs from across three distinct IVF batches and used the kinship analysis to further identify individual lines from within each batch that are derived from distinct sets of parents (**Fig. S4**).

**Figure 4.**
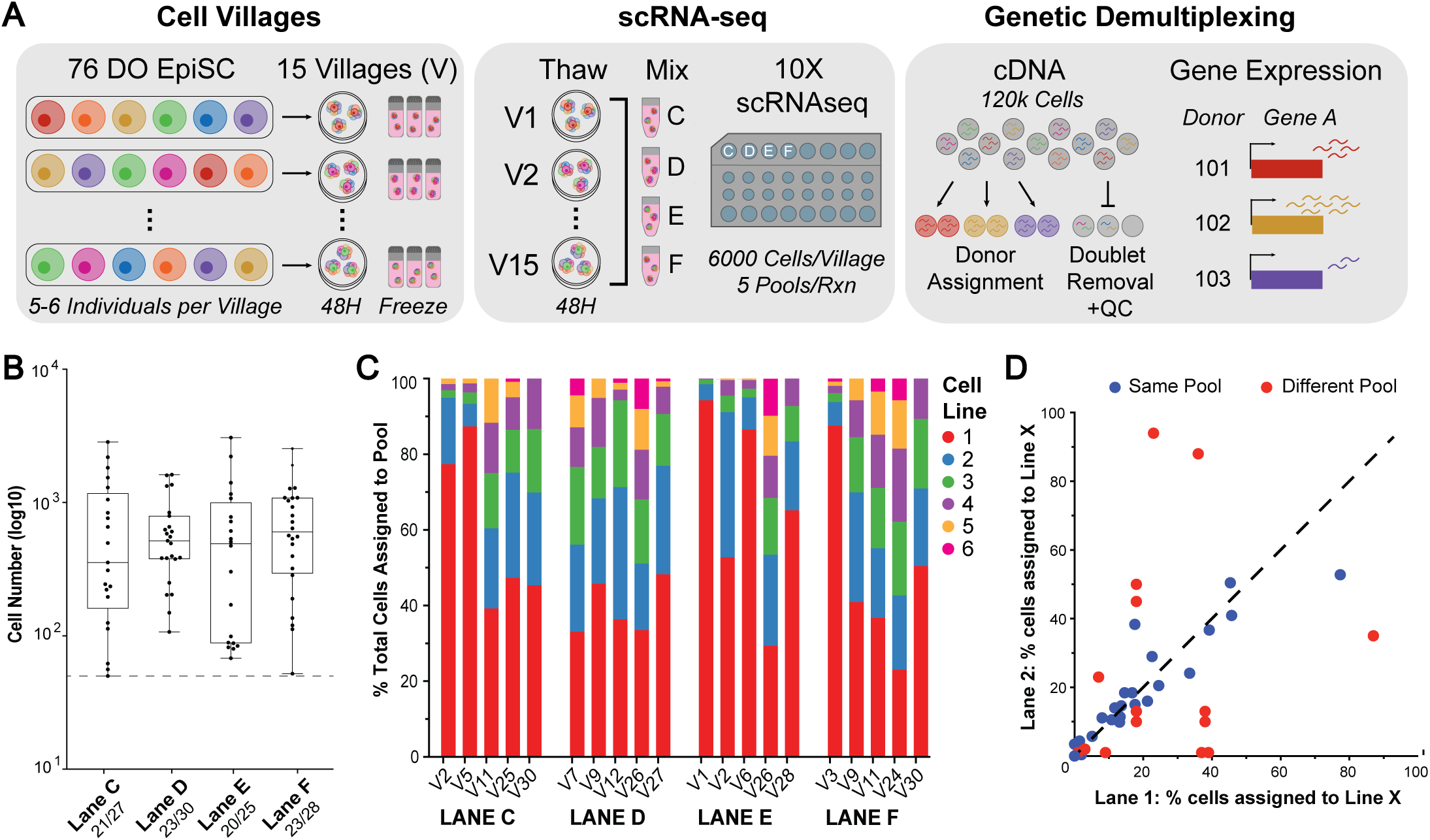
Multiplexed culture of DO EpiSCs in cell villages. **(A)** Experimental overview of cell villages strategy for pooled culture of DO EpiSC lines and genetic demultiplexing of scRNA-seq data. **(B)** Number of cells assigned to individual donors within each 10X Genomics sequencing channel following genetic demultiplexing and quality control analysis. EpiSC lines with >50 cells assigned to them are plotted as individual dots, and the number of individual lines that were recovered is indicated for each sample. **(C)** Relative abundance of each DO EpiSC line within a single cell village following genetic demultiplexing. Cell lines are indicated with numbers (1-6) based on their calculated relative abundance in each village (1 being the most abundant cell line, 6 being the least abundant cell line). **(D)** Relative contribution of the same cell line (line X) cultured in the same village and then included in a distinct sequencing channel (blue). This comparison was performed to assess how consistent the genetic demultiplexing results are across distinct lanes. In addition, we also determined the relative contribution of the same cell line (line X) within its village if it is included in two distinct cell villages (red). This comparison was performed to evaluate whether the specific cell lines included in each village of 5-6 lines impacts the relative growth of a given line within that village.

Briefly, individual EpiSC lines were cultured for 48 to 72 hours, dissociated into single cell suspension, and then 5-6 EpiSC lines were mixed in equal proportions into a single cell village (n = 15 total unique villages). Each village was then cultured for a single passage to generate a large stock for cryopreservation while also limiting the total time in pooled culture to minimize the potential for unequal growth/expansion of individual cell lines. Importantly, we also included a subset of the individual lines (n=33) in multiple villages to determine whether the behavior of a given cell line (i.e. its relative growth rate, transcriptome) are influenced by other individuals in a pool via cell competition or other cell nonautonomous variables^56^.

For large-scale characterization of the pools, we separately thawed and cultured 15 EpiSC villages, cultured them for 48 hours, harvested cells from each village, and combined 5 villages together (n = 28-30 total individuals) in equal proportion for single-cell RNA-sequencing [10X Genomics] (**Fig. 4A**; see methods). A subset of five villages were included in two separate sequencing lanes to assess the consistency of pooled culture composition. In total, we sequenced four sets of 28-30 DO EpiSC lines, loading ∼30,000 cells into each lane, and targeting a sequencing coverage of ∼40,000 reads per cell (**Fig. 4A**). To identify cells from specific individuals in the scRNA-seq datasets, we used Dropulations to perform genetic demultiplexing of single cells from within each village^61,62^. However, the SNP genotyping data from the GigaMUGA array is not sufficient for genetic demultiplexing using scRNA-seq data, as most SNPs assayed on the GigaMUGA array are located outside of genes, and thus would not be detectable in scRNA-seq reads^63^. To overcome this issue, we imputed the missing genotype information based on the haplotypes identified by the GigaMUGA markers (see methods)^64,65^.

Following genetic demultiplexing and quality control analysis of scRNA-seq data (see methods; **Figs. S5A-B**), we recovered 62,068 cells, representing 72 of the 76 input DO EpiSC lines. Among these 72 cell lines, 69 had >50 cells recovered, and these lines were used for downstream pseudo-bulk analyses. Using these data, we first assessed the number of cells assigned to each donor and village within each sequencing run to determine the variability of representation of each cell line. This revealed that several EpiSC lines had considerably increased or decreased cell numbers relative to other donors (**Fig. 4B**). To determine whether skewing of donor proportions occurred within the pooled culture rather than during processing of single cells for sequencing, we calculated the relative abundance of each cell line within its village (**Fig. 4C**). This analysis indicated moderate levels of donor skewing across most villages (e.g., the contribution of the most common donor is below 50%). More extensive skewing was detected in a smaller subset of the villages, in which a single donor is assigned to >75% of the total cells in the pool (e.g., V2), or in which some EpiSC lines were lost completely (V30), or both (V5). Analysis of the replicate pools included in more than one sequencing lane (but combined with distinct sets of villages for sequencing) indicated that the relative frequencies of each cell line in the cell village was consistent across sequencing lanes, indicating that the performance of the genetic demultiplexing pipeline was consistent across different sets of samples with different genotypes (**Figs. 4D, S5C**). For example, the relative contribution of each of the six donors in cell village 26 (V26) was relatively consistent when sequenced and demultiplexed across two 10X channels (**Fig. S5C**). In contrast, the set of EpiSCs that were included in multiple cell villages exhibited less consistency in their relative representation in distinct villages, which suggests that the relative abundance of each cell line within a pooled culture can be affected by both intrinsic and extrinsic factors (e.g., cell competition) (**Fig. 4D**).

Although we observed some variability in the relative representation of each EpiSC line within its village culture, we were nonetheless able to recover a sufficient number of EpiSCs (n ≥ 50 cells) for downstream pseudo-bulk gene expression analysis for 63/76 DO EpiSC lines included in the experiment. In summary, this pooled approach allowed for generation of single-cell transcriptional profiles for 63 DO EpiSC lines from just 15 individual cultures and four 10X sequencing lanes. This provides proof-of-concept data that this approach can be used for large-scale phenotyping of DO EpiSCs.

### Characterization of molecular and functional variation across DO EpiSCs

To perform additional characterization of DO EpiSC lines and to begin to assess phenotypic variation among the DO EpiSCs, we performed further analysis of genetically-demultiplexed scRNA-seq datasets. Clustering of merged scRNA-seq datasets revealed putative EpiSC clusters that express high levels of pluripotency markers (*Pou3f1*, *Pou5f1*, *Otx2)* as well as clusters expressing markers of primitive streak (*Foxa2*, *T*) and definitive endoderm (*Sox17*, *Cer1*) differentiation (**Fig. 5A-C**; see methods). The presence of primitive streak/DE cells in these cultures indicates that a subset of EpiSCs are undergoing spontaneous differentiation.

**Figure 5.**
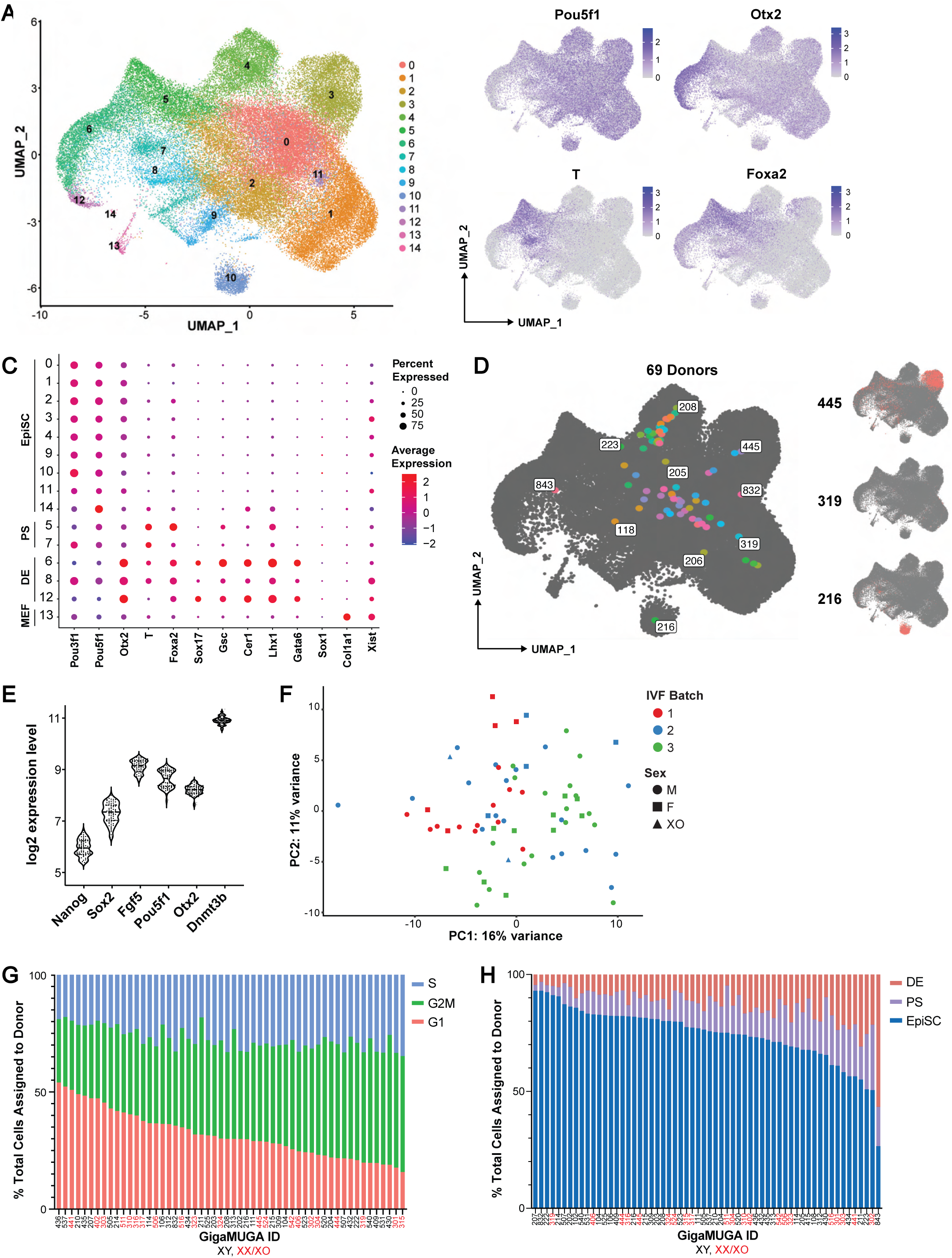
Mapping molecular and functional variation across DO EpiSCs. **(A)** UMAP representation of scRNA-seq data from 76 DO EpiSCs identifies 15 clusters from 62,068 cells. **(B)** Annotated UMAPs highlighting the expression of markers of pluripotency (*Pou5f1, Otx2)*, primitive streak (*T)*, and definitive endoderm (*Foxa2*, *Otx2*). *Otx2* is expressed in EpiSCs and DE. **(C)** Expression of selected genes within each of the assigned clusters. Clusters are grouped together into cell types based on marker gene expression. Cluster 13 appears to represent contaminating feeder cells (mouse embryonic fibroblasts) and was thus excluded from further analysis. **(D)** Distribution of cells from each EpiSC line across cell type clusters. The highlighted points represent the position of the average cell for each individual. For selected individuals (right panel), all cells assigned to that individual are highlighted on the UMAP. **(E)** Expression of selected primed pluripotency genes in individual pseudo-bulk expression data from merged EpiSC clusters (0, 1, 2, 3, 4, 9, 10, 11, 14). **(F)** PCA of pseudo-bulked gene expression data for cells assigned to each donor in putative EpiSC clusters (0, 1, 2, 3, 4, 9, 10, 11, 14). **(G)** Cell cycle profile of each DO EpiSC line inferred from scRNA-seq data. Male and female cell lines are color coded on the X-axis. **(H)** Relative proportions of cells from each DO EpiSC line that are assigned to pluripotent (EpiSC) versus spontaneously differentiated states (definitive endoderm [DE], primitive streak [PS]). Only cell lines with >50 total cells are shown.

To better understand whether heterogeneity within EpiSC clusters was driven by differences between EpiSC lines from different genetic backgrounds, we examined the distribution of each distinct EpiSC line across these clusters. Individual DO EpiSC lines exhibited different distributions across clusters, with some clusters consisting predominantly of cells from a single individual and others consisting of a mix of many different cell lines (**Fig. 5D**).

To evaluate gene expression in EpiSCs in greater depth, we generated pseudo-bulk transcriptomes of cells within EpiSC clusters for each individual genetic background. For these analyses, all cells within EpiSC clusters (0, 1, 2, 3, 4, 9, 10, 11, 14) from each individual were aggregated to produce a single EpiSC-pseudo- bulk transcriptome for each DO EpiSC line. Only samples with >50 EpiSC cells (n= 69) assigned were included in this analysis to ensure sufficient transcriptome coverage. Merged EpiSC transcriptomes for each individual exhibited strong expression of core pluripotency genes (*Nanog*, *Sox2*, *Fgf5*, *Pou5f1*, *Otx2, Dnmt3b*), indicating that genetically diverse DO EpiSCs can maintain gene expression signatures associated with primed pluripotency in cell villages (**Fig. 5E**). However, individual EpiSC lines do exhibit variation in the expression levels of some of these core regulators of pluripotency, consistent with transcriptional profiling data from naïve PSCs from DO mice (**Fig. 5E**)^25^. PCA of the pseudo-bulked transcriptomes indicates some clustering of donors by IVF batch or sex genotype (**Fig. 5F**). In addition to their transcriptional variation, we also observed significant variation in cell cycle dynamics across the EpiSC lines (**Fig. 5G)**. These differences could reflect intrinsic differences in the growth rate of each cell line, differences in propensity for cell cycle arrest, or competitive intercellular interactions between these lines that modulates their relative growth rates (i.e., cell competition).

Based on the observed differences in gene expression and cell cycle dynamics between EpiSCs, we evaluated whether these differences also manifest as changes in their propensity for spontaneous differentiation. For each EpiSC line, we determined the relative number of cells that were classified as undifferentiated EpiSCs vs. more differentiated progeny (primitive streak, definitive endoderm) (**Fig. 5H**). This revealed significant differences in the relative frequency of spontaneous differentiation across the DO EpiSC lines. Although a majority of the EpiSCs from different DO individuals maintained a high frequency of pluripotent EpiSCs, a small subset of the lines had a high frequency of spontaneous differentiation under these conditions.

Collectively, these data demonstrate that EpiSCs from genetically diverse DO mice can be stably cultured as cell villages. These proof-of-concept experiments indicate phenotypic variability among 69 genetically distinct individuals in gene expression, cell cycle dynamics, and differentiation propensity and highlight the utility of this platform for scalable culture and phenotyping of genetically diverse EpiSC lines and their differentiated progeny.

## DISCUSSION

Systematic studies in mouse and human have revealed pervasive genetic background effects on the phenotypic penetrance of many gene mutations *in vivo*^15,66–68^ and on the outcomes of *in vitro* directed differentiation^27,45,57,69^. These data highlight a critical challenge in modern developmental and stem cell biology -- understanding the contribution of genetic variation to quantitative phenotypic variation in developmental processes. To approach these complex questions, it will be necessary to develop new experimental approaches and model systems that can enable detailed mechanistic studies of molecular and cellular mechanisms and that incorporate natural genetic variation. Here, we sought to begin to address this important gap in the field by developing new experimental approaches and pluripotent stem cell resources from Diversity Outbred (DO) mice that have significant advantages compared to analogous human model systems.

First, we needed to improve standard mouse PSC culture to reduce technical variability in order to accurately measure the contribution of genetic variation to phenotypic variability. Current protocols require the addition of FGF/ERK and GSK3 inhibitors (2i) for the derivation and stable maintenance of mouse ESCs from the DO stock^52^. This is potentially problematic because ESCs cultured in these conditions for long periods are prone to genetic and epigenetic instability, especially in lines from females. This leads to increased phenotypic variation between lines that can complicate the subsequent directed differentiation of naïve cells^54,70–73^. Recently, our group and others have turned to primed PSC culture conditions (EpiSCs) that include inhibitors of canonical Wnt signaling pathway to culture mouse EpiSCs, which can potentially circumvent some of the aforementioned issues with naïve PSC culture and are a better starting point for directed differentiation to multiple lineages^46,74–76^. In addition, most human PSC studies use primed PSCs and so using EpiSCs allows for more direct comparisons between mouse and human differentiation models and thus increased synergy with available hPSC panels and differentiation methods. Here, we establish that these EpiSC culture conditions enable the stable culture of cells from diverse inbred and outbred genetic backgrounds, including both male and female mice (**Fig. 1**). This establishes a universal culture condition that can be used for the establishment, maintenance, and differentiation of large panels of PSCs from the DO stock.

The establishment of EpiSC lines from the inbred strains used to generate the Diversity Outbred stock (DO founder strains) enabled us to perform a strain survey to robustly assess the degree transcriptomic, epigenomic, and functional variation present across the DO founder strains. Several recent studies have observed molecular differences associated with genetic background among ESCs from the DO stock^25,41^ and in the naïve to primed transition of related inbred mouse PSCs^42,77^. However, there has been no systematic assessment of the effect of genetic variation on mouse primed PSCs. We generated a catalog of gene expression and chromatin states across 7/8 DO founder strains, establishing a significant contribution of genetic variation to gene regulatory variation in primed PSCs (**Fig. 1**). These data will also serve as valuable resource for subsequent analysis of gene regulation in DO EpiSCs. We extend these findings by surveying the differentiated progeny of these EpiSCs and find a similar contribution of genetic background to gene regulatory variation in definitive endoderm (**Fig. 2**). Together, these data further implicate genetic variation as a key regulator of pluripotency and early lineage specification and highlight the utility of stem cell models to identify genetic loci responsible for cellular and molecular phenotypes in cell state transitions.

The need for large panels of genetically diverse PSC lines from the DO stock represents a practical roadblock to the routine application of QTL studies in PSCs. For example, recent QTL mapping studies of ESCs from the DO stock used ∼200 unique ESC lines^25,41^. To overcome this issue, we implemented a pooled *in vitro* fertilization approach using multiple donors (10-15 males, 10-15 females) for each experiment, followed by direct derivation of ESCs from individual blastocyst stage embryos. This approach significantly reduces the need to house and breed DO mice, opening up the possibility for scalable derivation of genetically diverse ESCs from the DO stock. Using this approach, we were able to generate a panel of 230 new, genotyped ESC lines from the DO stock.

Culturing and differentiating large numbers of PSC lines also represents a logistical challenge for performing in vitro QTL studies. To address this issue, we employed a pooled cultured approach (“cell villages”) that has previously been used to phenotype large cohorts of human PSCs ^27,28^. We provide proof-of-concept data that pools of six genetically distinct DO EpiSC lines can be grown together, cryopreserved, and recovered for subsequent experiments with a low rate of drop out of individual cell lines. However, we did observe variation in the relative growth of individual cell lines within each pool, which can potentially complicate downstream applications (**Fig 4**). To characterize individual lines within pooled cultures, we performed scRNA-sequencing and adapted a genetic demultiplexing pipeline to assign single cells to specific DO genotypes (**Figs. 4-5**). These methods provide a foundation for future large-scale differentiation studies.

Our large stem cell bank, pooled phenotyping approach, and computational toolkit constitute a platform to perform forward genetic screens (QTL mapping) on large panels of PSCs from the DO stock and their differentiated progeny. Thus, our model serves as a foundation for studies seeking to dissect the mechanisms by which genetic variation influences cell fate specification and terminal differentiation during more complex developmental processes in mouse. In human, similar approaches have identified QTLs regulating cell type abundance in developing tissues and expression QTLs in developing cell types *in vitro* ^6,27,45,57^. In addition, findings from these studies should inform future studies seeking to generate cell types with higher purity and greater efficiency by identifying specific genetic backgrounds or variants associated with differentiation outcomes^69^. To enable such studies, our future work will include expanding the number of DO PSC lines in our bank, optimizing cell village culture of increased numbers of lines, and assessing other phenotyping approaches beyond scRNA-seq to identify other QTL types (chromatin accessibility^25^, protein abundance^41^, cell morphology^78^, cell type abundance QTLs following directed differentiation^27^). Following QTL mapping, candidate variants can potentially be further evaluated *in vivo* by generating PSC-derived mice and phenotyping the resulting embryos. In summary, our DO PSC resources and cellular model systems will allow for mapping of genotype-phenotype relationships and downstream mechanistic examination of developmental processes.

### Limitations of the study

One potential issue with employing mouse PSCs is that forward genetic screening may not identify variants directly relevant for human biology. Further, the DO founder strains encompass a larger range of genetic variation than what is found in human populations and are derived from inbred strains that have been recently subjected to strong, artificial selection for certain variants associated with domestication as laboratory strains^79^. While the underlying population genetics of human and mouse may differ, it is likely that mechanisms identified in the DO stock will converge on similar molecular and cellular pathways in human due to the high degree of conservation between the species^29,80^.

Although our current panel of 230 EpiSC lines is sufficiently large to map molecular traits with more simple genetic architecture, we are still likely underpowered for comprehensive mapping of QTLs for complex traits including *in vitro* differentiation efficiency. Similarly, we have only one ESC and EpiSC line for each genotype at present, and thus it can potentially be difficult to separate technical confounders from phenotypic differences caused by genetic variation. With an increasingly large panel of cell lines, pooled cell culture will become even more important to improve throughput and reduce technical variability. However, current levels of skewing in certain pools may limit the extent to which multiplexed experiments can be scaled, although we think it is likely that this issue can be mitigated by further optimization of experimental procedures for cell pooling and culture. Even though our current EpiSC conditions supported the culture of hundreds of DO lines, further optimization of mouse EpiSC culture conditions will be helpful to further reduce line-to-line variability and it will be beneficial in the long-term to develop robust, feeder-free culture conditions. In addition, it will be important to perform further in-depth studies of the genetic and epigenetic stability of EpiSC lines cultured in these conditions after extended *in vitro* passaging.

## ACKNOWLEDGEMENTS

This study would not have been possible without experimental support from the core facilities at MSKCC, including the Mouse Genetics Core Facility and the Integrated Genomics Operation Core. We would like to thank Steven Munger (The Jackson Laboratory), Christopher Baker (The Jackson Laboratory), Matthias Stadtfeld (Weill Cornell Medical College), Marty Yang (UCSF/Gladstone Institute), Lorenz Studer (MSKCC), and Anna Katerina-Hadjantonakis (MSKCC) for helpful discussions, advice, and/or comments on the manuscript throughout the course of this project. In addition, we would like to thank all members of the Vierbuchen Lab for their guidance, support, input, and advice. A/J iPSC lines were provided by Matthias Stadtfeld (WCMC).

## AUTHOR CONTRIBUTIONS

*Conceptualization*: R.G., E.K.C., T.V.

*Methodology*: R.G., E.K.C., S.D., T.V.

*Formal Analysis*: R.G., K.V., H.C., K.B., R.K., M.K.

*Investigation*: R.G., E.K.C., S.D., Y.F., J.L., L.S.

*Resources*: L.R., A.C.

*Data Curation*: R.G., K.G., M.K., R.K. *Writing – original draft*: R.G., T.V. *Writing – review and editing*: R.G., T.V. *Supervision*: T.V.

*Project Administration*: T.V.

*Funding Acquisition*: T.V.

## DATA AVAILABILITY

Source data and processed files are available in the Gene expression Omnibus under the accession numbers SUBMISSION PENDING (RNA and ATAC-seq datasets) and SUBMISSION PENDING (scRNA-seq).

## MATERIALS AND METHODS

### Inbred mESC derivation

The reference lines C57B6/J-44 and C57B6/J-51 were derived in the laboratory of Christopher Baker at the Jackson Laboratory as described in Czechanski et al. (2014)^52^. The remaining inbred mouse ESC lines (129S1/SvImJ, A/J, NOD/ShiLtJ, NZO/HiLtJ, CAST/EiJ, PWK/PhJ, WSB/EiJ) were generated in the laboratory of Laura Reinholdt at the Jackson Laboratory as used in Skelly et al. (2020)^25^. Early passage ESCs from all DO founder strains were frozen and shipped on dry ice to the Vierbuchen Lab in August 2020.

### Diversity Outbred mESC derivation

All the experiments involving live animals were performed by the Mouse Genetics Core Facility of Sloan Kettering Institute. Diversity Outbred (DO) mice were purchased from the Jackson Laboratory (J:DO, strain #009376). In-vitro fertilization (IVF) was performed essentially as described previously (Nakagata N. 2011. Meth Mol Biol 693:57-73.). Briefly, females were superovulated by administration of pregnant mere serum gonadotropin (PMSG, 5-7.5 IU, ProSpec Bio, catalog #HOR-272) approximately 62-64 hours prior to oocyte collection, followed by injection of human chorionic gonadotropin (HCG, 5-7.5 IU, Covetrus, catalog # 028938) 48 hours after PMSG. On the day of IVF, sperms of fertility-tested DO male mice were collected from the cauda epididymis and pre-incubated in a drop of sperm medium (TYH-MBCD medium (prepared in house) or Fertiup (CosmoBio, catalog #KYD-002-05-EX) for one hour. Cumulus masses containing oocytes were collected from the oviducts of superovulated females and placed in a drop of IVF medium (human tubal fluid (HTF) medium (prepared in house) or Cook IVF medium (Cook Medical, catalog # K-RVFE). One to 4 microliter of sperm was added to each IVF drop. In each batch of IVF, a non-sibling cohort of 10-15 males and 10-15 females was used. Sperms from 5 males were pooled into each of 4-6 sperm incubation drops in different combinations. Likewise, oocytes from 5 females were collected into each of 4-6 IVF drops, and then inseminated with distinct sperm pools. Three to 4 hours after insemination, eggs were rinsed to remove excess sperm and cumulus cells and cultured further. The following day, 2-cell stage embryos were collected and transferred to KSOM embryo culture medium (Embryo Max KSOM, Millipore-Sigma, catalog # MR-121-D), and further cultured for additional 2-3 days to hatching/hatched blastocysts. Blastocysts were then transferred to embryonic stem (ES) cell culture medium (Cellartis 3i mES/iPSC medium, Takara Bio, catalog # 1181722446) in a cell culture dish covered with mouse embryonic fibroblast (MEF) feeder cells. Subsequent steps of ES cell derivation were essentially based on methods previously described by Kiyonari et al. (Kiyonari H. et al., 2010. Genesis 48:317-327). Briefly, outgrowths of inner cell mass from seeded blastocysts cultured in 3i medium, typically for 1 week to 10 days, were manually picked. Cell clumps were dissociated with trypsin (0.25% Trypsin-EDTA, Gibco, catalog # 25200-056), and replated into MEF feeder-coated cell culture wells (typically, a 96-well plate). Cells were gradually expanded into larger cell culture wells, and the initial frozen stock vials were prepared from 6-well cell culture plates (typically, at passage # 3 to 4). A small number of remaining cells were re-plated onto gelatin-coated, feeder-free cell culture wells/dishes for further expansion and subsequent genomic DNA preparation with the Qiagen DNeasy Blood & Tissue Kit (#69504).

### Mouse ESC culture

Mouse PSC culture was performed as described in Medina-Cano et al. (2022)^46^. All mouse ESC lines were maintained on gelatin-coated dishes with irradiated mouse embryonic fibroblast feeder cells using serum/LIF media composed of DMEM (high glucose, GlutaMAX, HEPES), 1% nonessential amino acids, 1% sodium pyruvate, 1% penicillin-streptomycin, 0.1% 2-mercaptoethanol, 10% fetal bovine serum and 1000 units/ml ESGRO LIF. All ESC lines were cultured in serum/LIF media with 2i containing 3 µM CHIR99201 and 1 µM PD0325901. Media was changed daily, and ESCs were passaged upon 70% confluence at a 1:6 ratio using Accutase.

### Mouse EpiSC conversion and culture

ESC-to-EpiLC conversion was performed as described by Morgani et al. (2018)^81^. Briefly, ESC colonies were lifted using 0.5 µ/µl collagenase IV, centrifuged (200xg for 3 min at room temperature) and dissociated into a single-cell solution using Accutase. The ESC suspension was plated on fibronectin-coated plates (16.7 µg/ml) at a density of 17,500 cells/cm2 in N2B27 media supplemented with 12.5 ng/ml heat-stable recombinant human bFGF, 20 ng/ml activin A, and 1% Knockout Serum Replacement (Gibco, 1082028). N2B27 media consists of 50% DMEM-F12, 50% Neurobasal medium, 0.5% N2 supplement, 1% B27 supplement with vitamin A, 2 mM GlutaMAX, 1% penicillin-streptomycin and 0.1% 2-mercaptoethanol. The media was changed after 24 hours, and cells were converted to EpiSCs after 48 hours. For EpiLC-to-EpiSC conversion, EpiLCs were dissociated into small clumps (approximately three to five cells) with Accutase and plated at a density of ∼50,000-100,000 cells per cm^2^ on irradiated mouse fibroblast feeder cells in N2B27 with vitamin A supplemented with 20 ng/ml Activin A, 12.5 ng/ml heat-stable bFGF and Wnt inhibitor (175 nM NVP- TNKS656). EpiSC media was changed daily, and cells were passaged every ∼48 to 72 hours at a 1:6 ratio using 0.5 µ/µl collagenase IV followed by dissociation with Accutase into small clumps of three to five cells.

### DE directed differentiation

Directed differentiation of EpiSCs to DE progenitors was performed as described in Medina Cano et al. (2022)^46^. Briefly, EpiSCs were generally collected between four to eight passages in EpiSC conditions. For differentiations, EpiSC colonies were detached using collagenase IV (0.5 µ/μl), washed once with PBS, and then dissociated into a single-cell suspension using Accutase. EpiSCs were plated at a density of 110,000 cells/cm^2^ in chemically defined media (CDM) containing 50% IMDM, 50% Ham’s F12 Nutrient Mix with GlutaMAX, 1% chemically defined lipid concentrate, 450 µM monothioglycerol, 1% polyvinyl alcohol (w/v), 15 µg/ml Apo-transferrin, 0.5% GlutaMAX, 0.7 µg/ml insulin, and supplemented with 20 ng/ml activin A, 12.5 ng/ml heat-stable bFGF, 175 nM NVP-TNKS656, 1% Knockout Serum Replacement, and one of two Rho-associated kinase inhibitor cocktails – 2 µM thiazovivin, or 50 nm chroman and 5 µM emricasan. Plating media was removed 6 to 8 hours after seeding, cells were washed gently with PBS without calcium or magnesium (PBS-/-) and the media was changed to CDM supplemented with 3 µM CHIR99201 and 40 ng/ml Activin A. Sixteen hours after the first media change, cells were washed with PBS^-/-^ and the media was replaced with CDM supplemented with 100 ng/ml Activin A and 100 nM LDN-193189. After 24 h (46 total hours after cells were initially seeded), cells were fixed for immunostaining or collected for further analysis.

### Paraxial mesoderm directed differentiation

EpiSCs were collected between four to eight passages in EpiSC conditions. For differentiations, EpiSC colonies were first washed twice with PBS-/-, then detached using pre-warmed collagenase IV (0.5 µ/μl), and subsequently dissociated into a single-cell suspension using Accutase. EpiSCs were counted then plated at a density of 110,000 cells/cm^2^ in chemically defined media (CDM) containing insulin (0.7 ng/uL), FGF2 (12.5 ng/mL), Activin A (20 ng/mL), NVP-TNKS656 (175 nM), 1% Knockout Serum Replacement, Emricasan (5 uM), and Chroman 1 (50 nM). After 5-6 hours after seeding, cells were washed with PBS^-/-^ and the media was replaced with CDM containing 0.7 ng/uL insulin, supplemented with CHIRON (3 uM) and FGF2 (25 ng/uL). After 16 hours, the media was replenished. 24 hours later, the cells were fixed for immunostaining or collected for further analysis 46 total hours after the cells were initially seeded.

### Neurectoderm directed differentiation

EpiSCs were collected between four to eight passages in EpiSC conditions. For differentiations, EpiSC colonies were first washed twice with PBS-/-, then detached using pre-warmed collagenase IV (0.5 µ/μl), and subsequently dissociated into a single-cell suspension using Accutase. EpiSCs were counted then plated at a density of 110,000 cells/cm^2^ in Neurectoderm Differentiation Media containing N2B27 basal media (as described above, but with B27 supplement not containing vitamin A), 10 uM SB- 431542, 100 nM LDN- 193189. After 24 hours, the media was replenished. 24 hours later, the cells were fixed for immunostaining or collected for further analysis 48 total hours after the cells were initially seeded.

### Immunostaining

For immunostaining of adherent cells (EpiSCs and DE), cells were rinsed twice with PBS^-/-^ and fixed with 4% paraformaldehyde solution for 30 min at room temperature. After fixing, cells were washed twice with PBS^-/-^ and permeabilized with PBS^-/-^ containing 1% Triton X-100 (PBST) for 10 min at room temperature and then blocked for 30 min with PBS^-/-^ containing 0.3% Triton X-100 and 5% fetal bovine serum. Primary antibodies were diluted in PBST with 1% fetal bovine serum and incubated overnight at 4°C. Detailed information about the antibodies used can be found in Table S3. Following overnight incubation and three 10 min washes with PBST, cells were incubated with secondary antibodies for 2 hours at room temperature. Cells were rinsed with two 10 min washes of PBST before DAPI staining (1 µM) for 15 min at room temperature. Cells were rinsed with PBST and imaged using a Leica Dmi8 microscope.

### Flow Cytometry

Following DE differentiation, cells were dissociated in Accutase and washed in FACS buffer consisting of PBS-/-, 20% fetal bovine serum and 0.1% 0.5 M EDTA. CXCR4 antibody conjugated to Alexa Fluor 647 and live/dead zombie UV staining were diluted in FACS buffer at the manufacturer’s recommended concentration and incubated for 15 min at room temperature. Cells were washed and resuspended in FACS buffer with fixation/permeabilization diluent and fixation/permeabilization concentrate for 30 min at room temperature. The cells were then resuspended in permeabilization buffer with SOX17 antibody conjugated to Alexa Fluor 488 at the manufacturer’s recommended concentration for an additional 30 min at room temperature. Fixed cells were then analyzed using a Cytek Aurora cell sorter. Flow cytometry results were analyzed using FlowJo 10.10 software.

### MACS purification of DE

DE cells were isolated for ATAC-seq and RNA-seq analyses using MACS following the manufacturer’s instructions, with slight modifications. Cells were dissociated in Accutase, washed in FACS buffer, counted, and resuspended with rat anti-mouse CD184/CXCR4 Alexa Fluor 647-conjugated antibody at a concentration of 1 µl/106 cells and incubated for 1 hour at 4°C. Cells were then washed with FACS buffer, centrifuged (200 g for 3 min at 4°C) and resuspended in a solution composed by 80% FACS buffer and 20% anti-rat IgG MicroBeads (Miltenyi Biotec, 130-048-501). Cells were incubated in the mixture for 15 min at 4°C, then washed and resuspended in 500 µl buffer. The MS column was placed in an OctoMACS separator (Miltenyi Biotec, 130-048-501) on the MACS multistand. The cell suspension was added to the column and rinsed three times with buffer. Finally, the column was removed from the separator and the magnetically labeled cells were collected.

### RNA-sequencing sample collection and extraction

EpiSCs were collected using the collagenase and Accutase method as described above and DE cells were collected following MACS purification described above. Cells were counted and 5e6 to 1e6 cells were pelleted in PBS-/- and immediately resuspended in 1 mL of TRIzol (TRI Reagent) for storage at -80 degrees C. Phase separation in cells lysed in TRIzol was induced with 50% isopropanol and 0.5% 2-mercaptoethanol and RNA was extracted from the aqueous phase using the MagMAX mirVana Total RNA Isolation Kit (Thermo Fisher, A27828) on a KingFisher Flex Magnetic Particle Processor according to the manufacturer’s protocol with 1-2 million cells input. Samples were eluted in 38 µl elution buffer.

### Transcriptome sequencing of EpiSC samples

After RiboGreen quantification and quality control using an Agilent BioAnalyzer, 100-500 ng of total RNA with RIN values of 9.5-10 underwent polyA selection and TruSeq library preparation according to instructions provided by Illumina (TruSeq Stranded mRNA LT Kit), with eight cycles of PCR. Samples were barcoded and run on a NovaSeq 6000 in a PE100 run, using the NovaSeq 6000 S4 Reagent Kit (200 Cycles) (Illumina) at the Integrated Genomics Operation at SKI.

### Transcriptome sequencing of DE samples

After RiboGreen quantification and quality control using an Agilent BioAnalyzer, 2 ng total RNA with RNA integrity numbers ranging from 7.7 to 10 underwent amplification using the SMART-Seq v4 Ultra Low Input RNA Kit, with 12 cycles of amplification. Subsequently, 10 ng of amplified cDNA was used to prepare libraries with the KAPA Hyper Prep Kit using eight cycles of PCR. Samples were barcoded and run on a NovaSeq 6000 in a PE100 run, using the NovaSeq 6000 S4 Reagent Kit (200 Cycles) (Illumina) at the Integrated Genomics Operation at SKI.

### ATAC-sequencing

ATAC-seq was performed as previously described (Buenrostro et al., 2015; see also dx.doi.org/10.17504/protocols.io.bv9mn946) with minor modifications. Briefly, cells were dissociated into single cells, filtered through a Flowmi cell strainer and the nuclei were isolated by incubation with lysis buffer [10 mM Tris-HCl pH 7.4, 10 mM NaCl, 3 mM MgCl2, 0.1% Tween-20, 0.1% NP-40, 0.01% digitonin and 1% bovine serum albumin (BSA)] for 5 min at 4°C. 20,000 and 40,000 nuclei were isolated per sample, and the DNA was tagmented with Tn5 (Illumina) and amplified using NEBNext® High-Fidelity 2X PCR Master Mix (NEB). The number of cycles was estimated by qPCR. DNA tagmentation efficacy was evaluated with a Bioanalyzer 2100 (Agilent Technologies) and the DNA amounts calculated with Qubit. i5 and i7 primer sequences were obtained from Mezger et al. (2018). The resulting DNA libraries were sequenced using the NextSeq550 system (Illumina) at the Integrated Genomics Operation and Center for Epigenetics Research at SKI.

### RNA-sequencing analysis

RNA sequencing reads were 3’ trimmed for base quality 15 and adapter sequences using version 0.4.5 of TrimGalore (https://www.bioinformatics.babraham.ac.uk/projects/trim_galore), and then aligned to mouse assembly mm10 with STAR v2.7.10b using default parameters. Data quality and transcript coverage were assessed using the tool CollectRNASeqMetrics from Picard (http://broadinstitute.github.io/picard/). Read count tables were generated with HTSeq v0.9.1. Normalization and expression dynamics were evaluated with DESeq2 using the default parameters and outliers were assessed by sample grouping in principal component analysis. Gene set enrichment analysis (GSEA, http://software.broadinstitute.org/gsea) was run against MSigDB v6 using the pre-ranked option and log2 fold change for pairwise comparisons. Kmeans clustering was performed on the normalized counts from the superset of all differentially expressed genes in R using the function pheatmap.

### ATAC-seq analysis

ATAC sequencing reads were trimmed and filtered for quality threshold 15 and adapter content using version 0.4.5 of TrimGalore (https://www.bioinformatics.babraham.ac.uk/projects/trim_galore) and using cutadapt (v1.15) and FastQC (v0.11.5). Reads were aligned to mouse assembly mm10 with version 2.3.4.1 of bowtie2 (http://bowtie-bio.sourceforge.net/bowtie2/index.shtml) and were deduplicated using MarkDuplicates in version 2.16.0 of Picard Tools. To ascertain regions of chromatin accessibility, MACS2 (https://github.com/taoliu/MACS) was used with a p-value setting of 0.001. The BEDTools suite v2.29.2 (http://bedtools.readthedocs.io) was used to create read density profiles normalized to ten million uniquely mapped reads. A global peak atlas was created by first removing blacklisted regions (http://mitra.stanford.edu/kundaje/akundaje/release/blacklists/ mm10-mouse/mm10.blacklist.bed.gz) then taking 500 bp windows around peak summits and counting reads with featureCounts (v1.6.1). DESeq2 was used to calculate differential enrichment for all pairwise contrasts. Peak-gene associations were created by assigning all intragenic peaks to that gene, and otherwise using linear genomic distance to transcription start site. Peak intersections were calculated using bedtools and intersectBed with 1 bp overlap. Gene set enrichment analysis (GSEA, http://software.broadinstitute.org/gsea) was performed with the pre-ranked option and default parameters, where each gene was assigned the single peak with the largest (in magnitude) log2 fold change associated with it. Motif signatures were obtained using Homer v4.5 (http://homer.ucsd.edu).

Profile and tornado plots were created using deepTools v3.3.0 by running computeMatrix and plotHeatmap on normalized bigwigs with average signal sampled in 25 bp windows and flanking region defined by the surrounding 3 kb. Kmeans clustering was performed on the normalized counts from the superset of all differentially accessible regions in R using the function pheatmap.

### Genotyping of DO PSCs

For other lines, a cell pellet of 5e5 to 1e6 was collected from ESCs grown on feeders for 1-2 passages. DNA was extracted using the Qiagen DNeasy Blood & Tissue Kit followed by the Zymo DNA Clean and Concentrator. Double-stranded DNA content was measured using a NanoDrop OneC Spectrophotometer (Thermo Fisher, #13-400-518). Extracted DNA was aliquoted in a 96-well qPCR plate in at least 10 uL of volume at greater than 20 ng/uL concentration in groups of at least 24 samples. Samples were shipped to Neogen Genomics (Lincoln, NE) on dry ice overnight for sequencing via the GigaMUGA genotyping array on the Illumina Infinium platform. The array was optimized for the DO population and covers 143,000+ SNPS with an average spacing of 22.5 kB. The sample sequences were processed using the R/qtl2 package to encode the SNP genotypes as described in Broman et al. (2019)^82^. Subsequently, we performed a kinship analysis to compare the genotype probabilities between samples and define the degree of relatedness between lines from the same IVF preparation as adapted from Broman and Sen (2009) at https://smcclatchy.github.io/mapping/.

### Quality Control of the Diversity Outbred Mice Genotypes

Using the R/qtl2 package (https://github.com/rqtl/qtl2), the following metrics were computed for each diversity outbred mouse: X chromosome heterozygosity, number of crossovers, proportion missing data, and proportion heterozygous sites. The Y chromosome intensity per mouse by averaging the average Y chromosome microarray signal across all SNPs. Mice genotypes were filtered using the following criteria: either have X chromosome heterozygosity < 0.1 and Y chromosome intensity > 0.2 (male) or have X chromosome heterozygosity > 0.1 and Y chromosome intensity < 0.2 (female), must have between 600 and 900 crossovers, must have less than or equal to 2% proportion missing genotype, must have proportion heterozygosity between 0.75 and 0.9. Plots of the B allele frequency and the log R ratio per sample were generated using the karyoploteR package (https://github.com/bernatgel/karyoploteR).

### PCA Visualization of Diversity Outbred Mouse Genotypes

To reduce the dimensionality of the genotype data, principal component analysis was performed on the autosomal genotypes of the 230 diversity outbred mice that passed our defined QC metrics. From the GigaMUGA array data, ‘A’ was replaced with 0, ‘H’ was replaced with 1, and ‘B’ was replaced with 2 to represent frequency of the minor allele (cite GigaMUGA?). Missing genotypes for each SNP indicated by ‘-‘ were imputed as 0 for the purpose of running the principal component analysis only. SNPs where all samples had the same genotype were removed, resulting in 108,578 SNPs total. PCA was performed using the prcomp() function in R. The genotypes were scaled to have unit variance prior to PCA using the ‘scale = TRUE’ argument.

### Cell pooling

DO EpiSC lines were thawed onto irradiated MEFs in EpiSC media. Cells were cultured after thaw for 2-4 days until 50-70% confluent at which time colonies were lifted using collagenase, further dissociated to single cells using Accutase, and counted as described above. Primary DO pools were generated by combining equal numbers of 5-6 independent lines in 10 cm dishes (2e6 total cells per dish) on irradiated MEFs in EpiSC media. After 2-3 days, cells were treated with Accutase to collect EpiSCs and MEFs and frozen in basal N2B27 with 10% DMSO.

### Single-cell RNA-sequencing

EpiSCs were collected using collagenase and Accutase as per the EpiSC 48 hours post-thaw. The resulting cell suspension was used for 10x scRNA-seq (10x Genomics, Single Cell 3′ Kit v3.1, dual index) following the manufacturer’s directions. We combined four cell villages in equal number to a total of 30,000 cells per sequencing channel. We targeted a sequencing depth of 40,000 reads per cell (1.2e9 total reads per sequencing channel) for genetic demultiplexing.

### Demultiplexing scRNA-sequencing of DO cell villages

Complete SNP genotypes for each EpiSC line were imputed from GigaMUGA SNP genotyping data, as described in Lobo et al., (2021)^65^ and converted to .VCF files. Raw sequencing read data were processed using the Dropseq computational pipeline with default parameters^28,61,62^. Raw reads were aligned to the mouse reference genome GRCm38 (mm10), giving a matrix representing unique molecular identifers (UMIs) per cell barcode per gene. To demultiplex the cell village, we used the VCF files containing imputed genotype information as input for the Dropseq AssignCellsToSamples function. Based on the output from the DetectDoublets function, we developed a custom assignment method. By applying the Rosner Test for outliers and analyzing the absolute distances of previously calculated donor likelihoods, we categorized each cell as a singlet or doublet. Only singlets were utilized for downstream analysis. The raw UMI matrices for each sample were processed using R v4.3.1 with the Seurat package v5.0.1. Matrices were filtered to remove cells with less than 200 genes detected and cells with >10% of reads mapping to mitochondrial RNA. Data from all samples was then integrated to combine and account for batch effects using the IntegrateLayers function using CCAIntegration as integration method.

### Differential gene expression analysis

The top 2000 most variably expressed genes were used for dimensionality reduction, firstly by principal component analysis (PCA) and subsequently by uniform manifold projection (UMAP), selecting PCs 1:20 that explained the majority of the variance observed (assessed by elbow plots). A shared nearest-neighbour graph was constructed in PCA-space using PCs 1:20 with Seurat FindNeighbors function. Clusters are identifed within this graph using Seurat FindClusters function, optimizing the modularity with the Louvain algorithm. The resolution parameter to control cluster granularity was manually selected at 0.5. Cluster marker genes were identifed with FindAllMarkers function only testing genes that are detected in a minimum fraction of 0.1 cells and using a logFC cutoff of 0.1. Pseudo-bulk transcriptomes were generated using Seurat AggregateExpression function and counts were log-normalized before downstream analysis. Limma removeBatchEffect effect was utilized to account for lane dependent batch effects. Genes differentially expressed between groups were identifed per cluster using Seurats fndMarkers utilizing Wilcoxon Rank Sum test with default parameters.

## SUPPLEMENTARY FIGURE LEGENDS

**Supplementary Figure 1.**
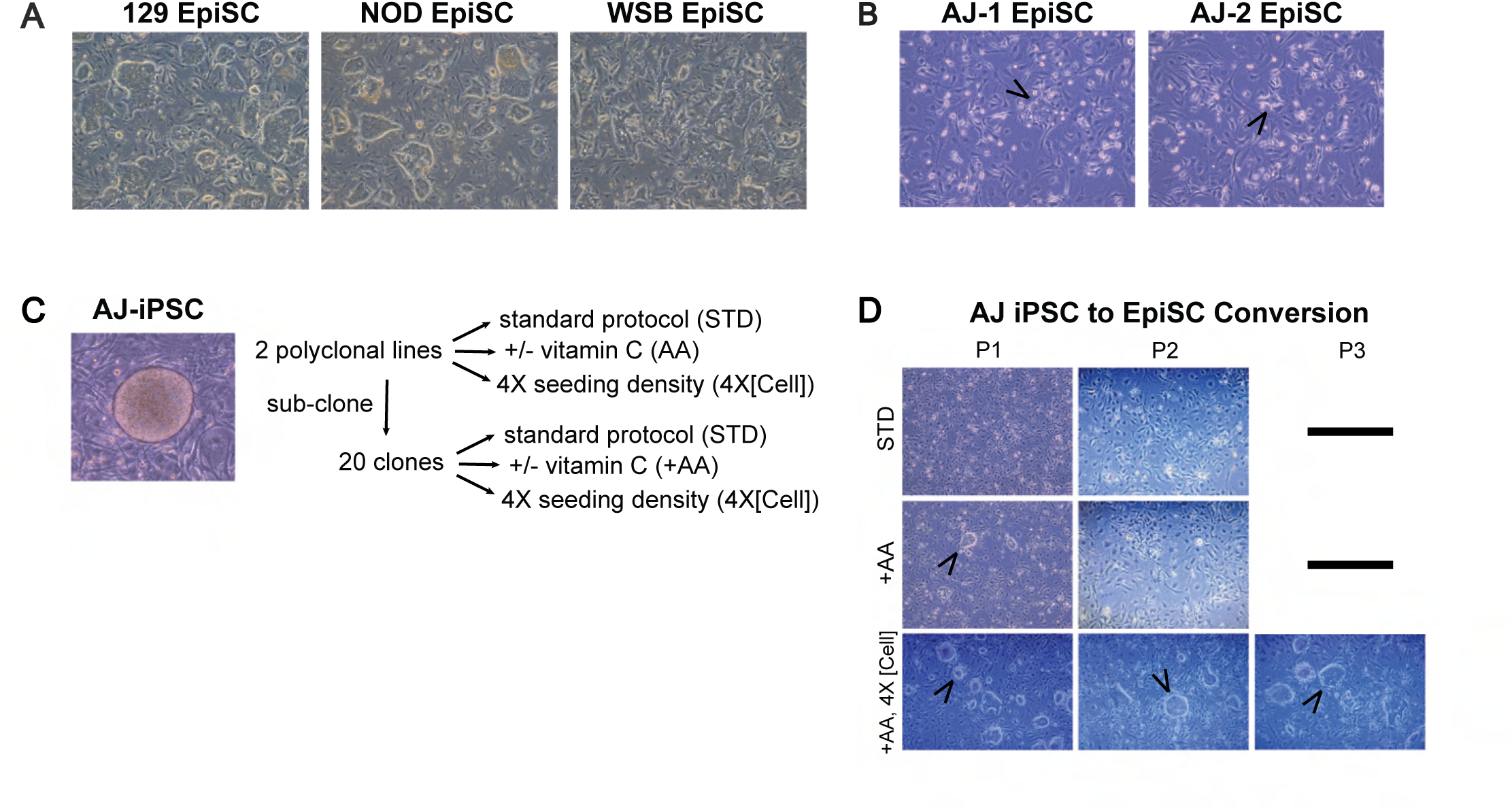
Additional characterization of EpiSCs from DO founder strains. **(A)** Representative phase contrast images of three EpiSCs from the indicated DO founder strains. **(B)** Phase contrast images of nascent AJ EpiSCs after 48 hours in EpiSC culture media. Arrowheads indicate EpiSC colonies. **(C)** Outline of additional iPSC to EpiSC conversion experiments. AJ iPSC lines (polyclonal population, or isolated subclones) were used for EpiSC conversion. Alternate conditions for EpiSC conversion were evaluated to try and improve viability of AJ EpiSCs, including addition of vitamin C (AA) or increased seeding density (all conditions cultured on irradiated MEFs). **(D)** Phase contrast images of AJ iPSC-derived EpiSC lines following 48-hour EpiLC conversion over three subsequent passages in standard EpiSC media (STD) conditions, with vitamin C media supplement (+AA), or at a higher EpiLC seeding concentration (+AA, [4X cell]). Conditions with dashed lines no longer contained observable EpiSC colonies.

**Supplementary Figure 2.**
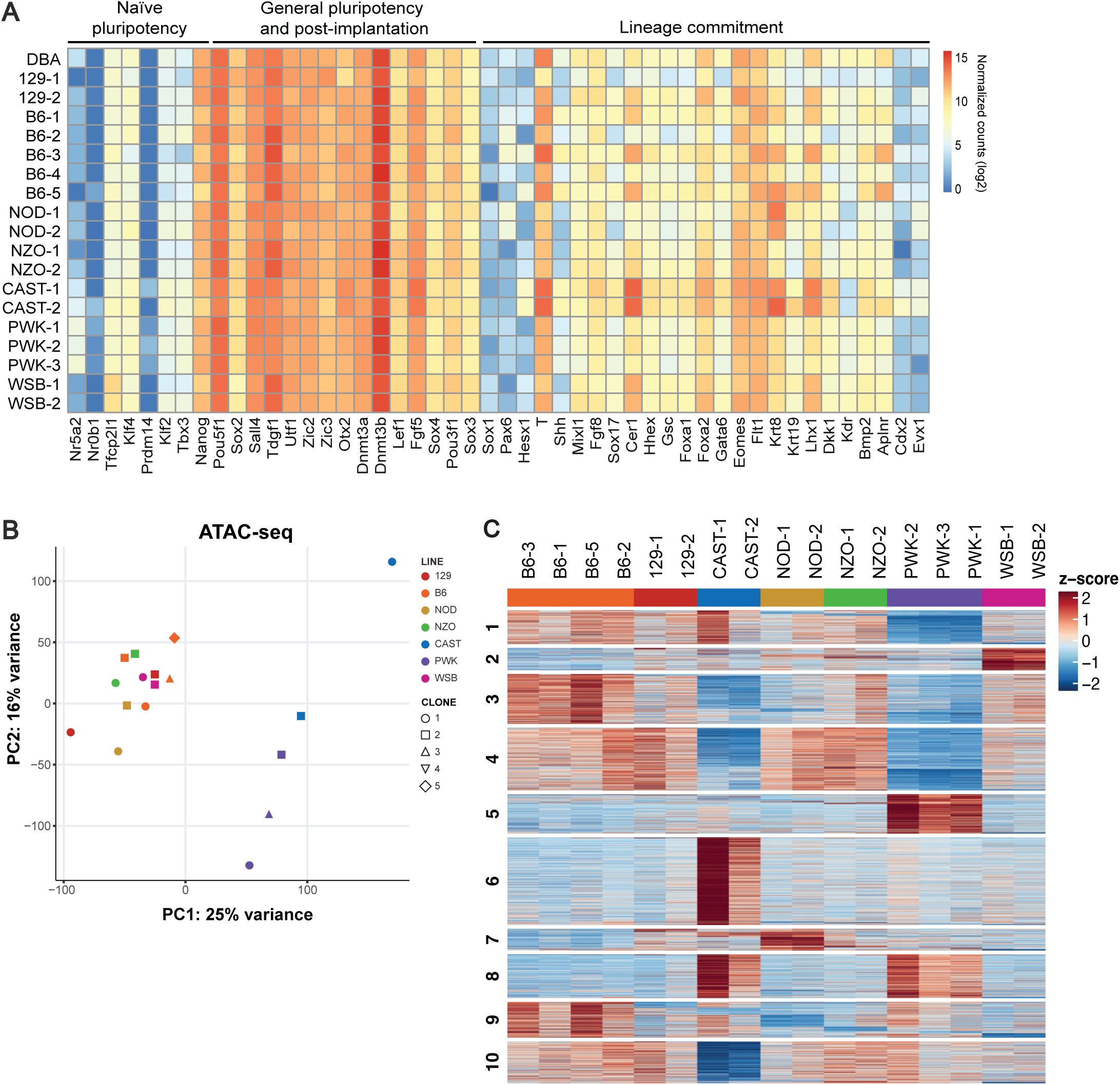
Additional EpiSC RNA-sequencing and ATAC-sequencing results from DO founder strain survey. **(A)** Heatmap of bulk RNA-seq data for selected genes associated with naïve pluripotency, post-implantation epiblast, or differentiation into specific germ layers (lineage commitment). **(B)** PCA of chromatin accessibility (ATAC-seq) in EpiSCs from DO founder strains. DBA/2J EpiSCs are included for comparison. **(C)** k-means clustering (k =10) of bulk chromatin accessibility differences across DO founder EpiSCs.

**Supplementary Figure 3.**
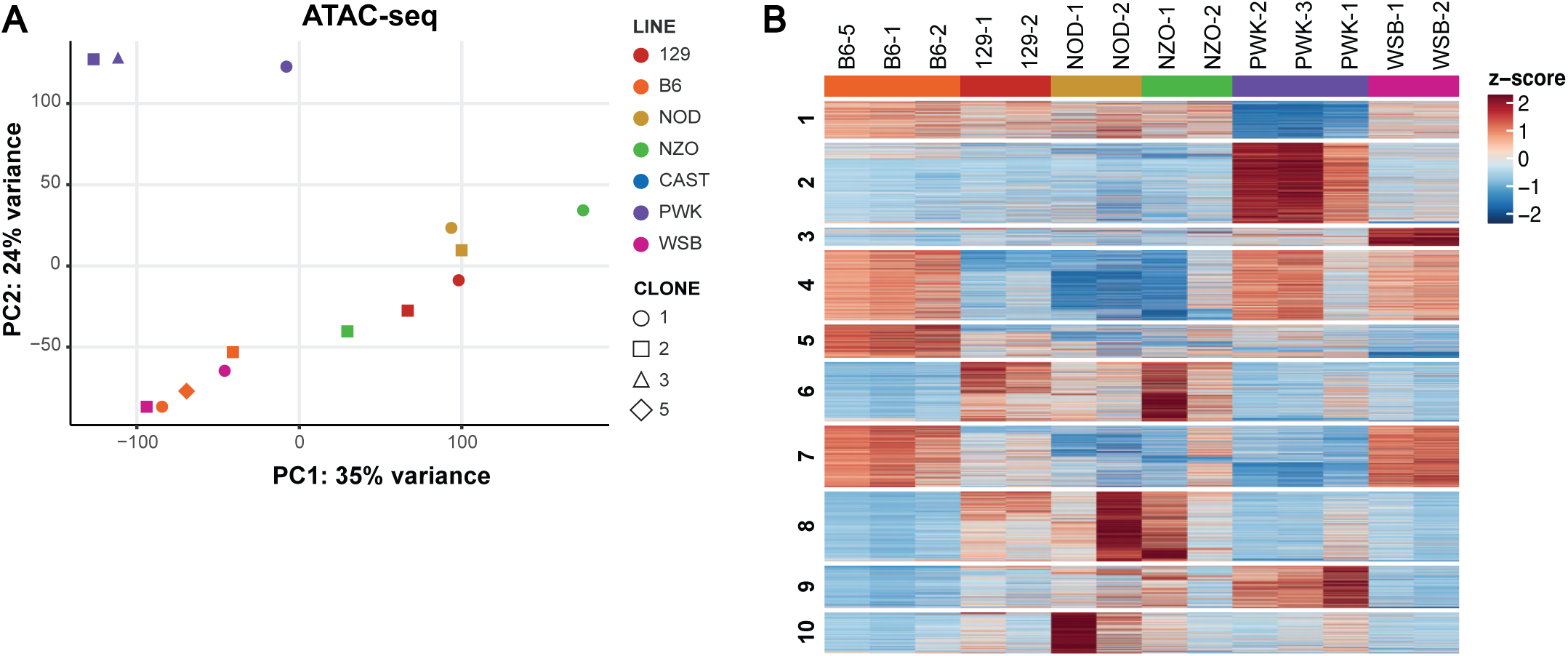
Summary of ATAC-seq data from definitive endoderm from DO founder strains. **(A)** PCA of chromatin accessibility (ATAC-seq) in purified DE from each of the DO founder strains. DBA/2J DE is included for comparison. **(B)** k-means clustering (k =10) of bulk chromatin accessibility differences of purified DE from DO founder strains.

**Supplementary Figure 4.**
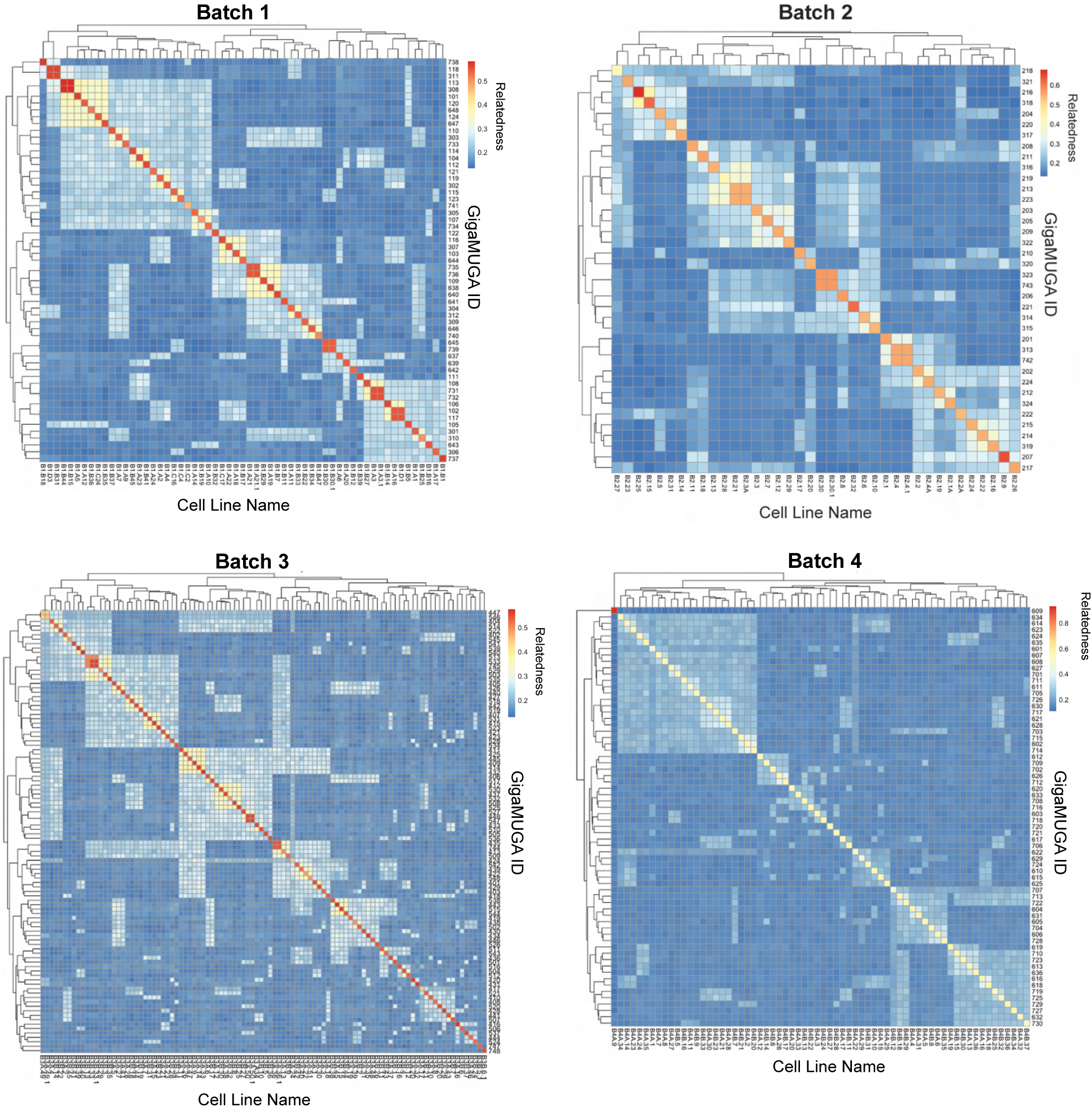
Pooled *in vitro* fertilization generates genetically diverse EpiSC lines with distinct parentage. Kinship matrices describing the degree of genetic similarity (relatedness) between DO PSCs based on GigaMUGA SNP genotypes. Results from each of the 4 pooled IVF batches are shown.

**Supplementary Figure 5.**
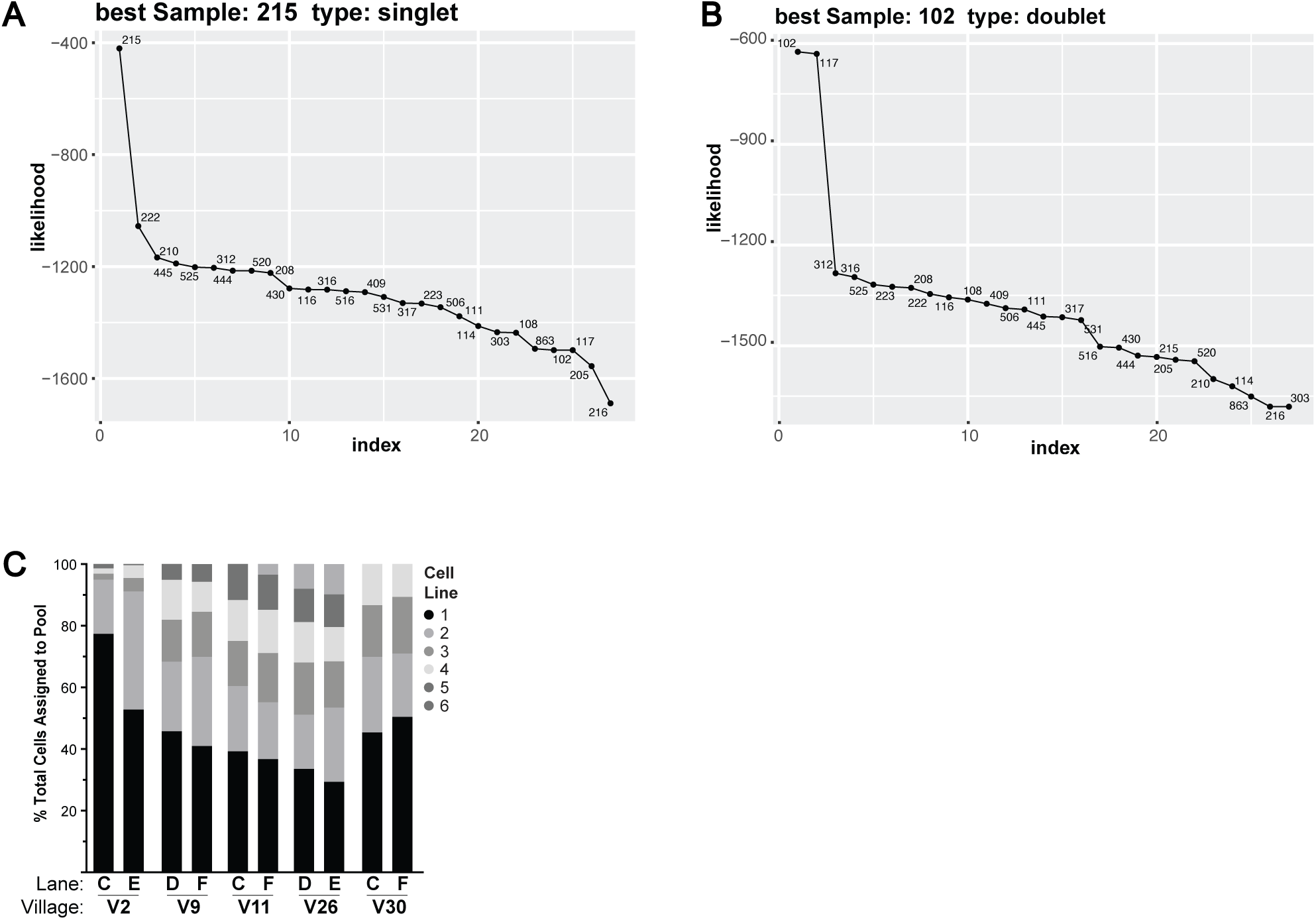
Additional information related to cell village culture experiments. **(A-B)** Example outputs from Dropulations demultiplexing pipeline ranking the likelihood of individual donor genotypes being assigned to a given cell based on SNP genotypes. Dropulations output indicating assignment of a specific cell to one individual donor (singlet) (A) or indicative of a doublet (i.e., two donor genotypes identified) (B). **(C)** Proportion of cells assigned to donor genotype within the same pool sequenced across multiple 10X lanes. This comparison was performed to assess whether the relative abundance of EpiSC lines within the same pool can be consistently detected across different 10X sequencing lanes.

## SUPPLEMENTARY TABLES

**Supplementary Table 1: Diversity Outbred founder stem cell lines.**

**Supplementary Table 2: Diversity Outbred stem cell lines.**

**Supplementary Table 3: List of antibodies used. Supplementary Table 4: List of reagents used.**

## REFERENCES

1. Uffelmann, E. et al. Genome-wide association studies. Nat Rev Methods Primers 1, 1–21 (2021).

2. Visscher, P. M. et al. 10 Years of GWAS Discovery: Biology, Function, and Translation. The American Journal of Human Genetics 101, 5–22 (2017).

3. Umans, B. D., Battle, A. & Gilad, Y. Where are the disease-associated eQTLs? Trends Genet 37, 109–124 (2021).

4. Claussnitzer, M. et al. FTO Obesity Variant Circuitry and Adipocyte Browning in Humans. N Engl J Med 373, 895–907 (2015).

5. D’Antonio, M. et al. Fine mapping spatiotemporal mechanisms of genetic variants underlying cardiac traits and disease. Nat Commun 14, 1132 (2023).

6. Strober, B. J. et al. Dynamic genetic regulation of gene expression during cellular differentiation. Science 364, 1287–1290 (2019).

7. Walker, R. L. et al. Genetic Control of Expression and Splicing in Developing Human Brain Informs Disease Mechanisms. Cell 179, 750–771.e22 (2019).

8. Aygün, N. et al. Brain-trait-associated variants impact cell-type-specific gene regulation during neurogenesis. Am J Hum Genet 108, 1647–1668 (2021).

9. Gasch, A. P., Payseur, B. A. & Pool, J. E. The Power of Natural Variation for Model Organism Biology. Trends Genet 32, 147–154 (2016).

10. Ghosh, S., Nehme, R. & Barrett, L. E. Greater genetic diversity is needed in human pluripotent stem cell models. Nat Commun 13, 7301 (2022).

11. Chandler, C. H., Chari, S. & Dworkin, I. Does your gene need a background check? How genetic background impacts the analysis of mutations, genes, and evolution. Trends Genet 29, 358–366 (2013).

12. Horwitz, R., Riley, E. A. U., Millan, M. T. & Gunawardane, R. N. It’s time to incorporate diversity into our basic science and disease models. Nat Cell Biol 23, 1213–1214 (2021).

13. Voelkl, B. et al. Reproducibility of animal research in light of biological variation. Nat Rev Neurosci 21, 384– 393 (2020).

14. Tabbaa, M., Knoll, A. & Levitt, P. Mouse population genetics phenocopies heterogeneity of human *Chd8* haploinsufficiency. Neuron 111, 539–556.e5 (2023).

15. Sittig, L. J. et al. Genetic Background Limits Generalizability of Genotype-Phenotype Relationships. Neuron 91, 1253–1259 (2016).

16. Neuner, S. M., Heuer, S. E., Huentelman, M. J., O’Connell, K. M. S. & Kaczorowski, C. C. Harnessing Genetic Complexity to Enhance Translatability of Alzheimer’s Disease Mouse Models: A Path toward Precision Medicine. Neuron 101, 399–411.e5 (2019).

17. Farbehi, N. et al. Integrating population genetics, stem cell biology and cellular genomics to study complex human diseases. Nat Genet 56, 758–766 (2024).

18. Warren, C. R. & Cowan, C. A. Humanity in a Dish: Population Genetics with iPSCs. Trends in Cell Biology 28, 46–57 (2018).

19. Swanzey, E., O’Connor, C. & Reinholdt, L. G. Mouse Genetic Reference Populations: Cellular Platforms for Integrative Systems Genetics. Trends in Genetics 0, (2020).

20. Saul, M. C., Philip, V. M., Reinholdt, L. G. & Chesler, E. J. High-Diversity Mouse Populations for Complex Traits. Trends in Genetics 35, 501–514 (2019).

21. Pashos, E. E. et al. Large, Diverse Population Cohorts of hiPSCs and Derived Hepatocyte-like Cells Reveal Functional Genetic Variation at Blood Lipid-Associated Loci. Cell Stem Cell 20, 558–570.e10 (2017).

22. Warren, C. R. et al. Induced Pluripotent Stem Cell Differentiation Enables Functional Validation of GWAS Variants in Metabolic Disease. Cell Stem Cell 20, 547–557.e7 (2017).

23. DeBoever, C. et al. Large-Scale Profiling Reveals the Influence of Genetic Variation on Gene Expression in Human Induced Pluripotent Stem Cells. Cell Stem Cell 20, 533–546.e7 (2017).

24. Kilpinen, H. et al. Common genetic variation drives molecular heterogeneity in human iPSCs. Nature 546, 370–375 (2017).

25. Skelly, D. A. et al. Mapping the Effects of Genetic Variation on Chromatin State and Gene Expression Reveals Loci That Control Ground State Pluripotency. Cell Stem Cell 27, 459–469.e8 (2020).

26. van der Wijst, M. et al. The single-cell eQTLGen consortium. eLife 9, e52155 (2020).

27. Cuomo, A. S. E. et al. Single-cell RNA-sequencing of differentiating iPS cells reveals dynamic genetic effects on gene expression. Nature Communications 11, 810 (2020).

28. Wells, M. F. et al. Natural variation in gene expression and viral susceptibility revealed by neural progenitor cell villages. Cell Stem Cell (2023) doi:10.1016/j.stem.2023.01.010.

29. Allayee, H. et al. Systems genetics approaches for understanding complex traits with relevance for human disease. Elife 12, e91004 (2023).

30. Fu, J., Warmflash, A. & Lutolf, M. P. Stem-cell-based embryo models for fundamental research and translation. Nature Materials 1–13 (2020) doi:10.1038/s41563-020-00829-9.

31. Tewary, M., Shakiba, N. & Zandstra, P. W. Stem cell bioengineering: building from stem cell biology. Nature Reviews Genetics 19, 595–614 (2018).

32. Churchill, G. A., Gatti, D. M., Munger, S. C. & Svenson, K. L. The Diversity Outbred mouse population. Mamm Genome 23, 713–718 (2012).

33. Roberts, A., Pardo-Manuel de Villena, F., Wang, W., McMillan, L. & Threadgill, D. W. The polymorphism architecture of mouse genetic resources elucidated using genome-wide resequencing data: implications for QTL discovery and systems genetics. Mamm Genome 18, 473–481 (2007).

34. Svenson, K. L. et al. High-Resolution Genetic Mapping Using the Mouse Diversity Outbred Population. Genetics 190, 437–447 (2012).

35. Logan, R. W. et al. High-precision genetic mapping of behavioral traits in the diversity outbred mouse population. Genes Brain Behav 12, 424–437 (2013).

36. Hsiao, K. et al. A Thalamic Orphan Receptor Drives Variability in Short-Term Memory. Cell 183, 522–536.e19 (2020).

37. Winter, J. M. et al. Mapping Complex Traits in a Diversity Outbred F1 Mouse Population Identifies Germline Modifiers of Metastasis in Human Prostate Cancer. cels 4, 31–45.e6 (2017).

38. French, J. E. et al. Diversity Outbred Mice Identify Population-Based Exposure Thresholds and Genetic Factors that Influence Benzene-Induced Genotoxicity. Environmental Health Perspectives 123, 237–245 (2015).

39. Keller, M. P. et al. Genetic Drivers of Pancreatic Islet Function. Genetics 209, 335–356 (2018).

40. Hackett, S. R. et al. The Molecular Architecture of Variable Lifespan in Diversity Outbred Mice. 2023.10.26.564069 Preprint at 10.1101/2023.10.26.564069 (2023).

41. Aydin, S. et al. Genetic dissection of the pluripotent proteome through multi-omics data integration. 2022.04.22.489216 Preprint at 10.1101/2022.04.22.489216 (2022).

42. Ortmann, D. et al. Naive Pluripotent Stem Cells Exhibit Phenotypic Variability that Is Driven by Genetic Variation. Cell Stem Cell 27, 470–481.e6 (2020).

43. Byers, C., et al. Genetic Control of Pluripotency Epigenome Determines Differentiation Bias in Mouse Embryonic Stem Cells. http://biorxiv.org/lookup/doi/10.1101/2021.01.15.426861 (2021) doi:10.1101/2021.01.15.426861.

44. Jerber, J. et al. Population-scale single-cell RNA-seq profiling across dopaminergic neuron differentiation. bioRxiv 2020.05.21.103820 (2020) doi:10.1101/2020.05.21.103820.

45. Cederquist, G. Y. et al. A Multiplex Human Pluripotent Stem Cell Platform Defines Molecular and Functional Subclasses of Autism-Related Genes. Cell Stem Cell 27, 35–49.e6 (2020).

46. Medina-Cano, D. et al. Rapid and robust directed differentiation of mouse epiblast stem cells into definitive endoderm and forebrain organoids. Development 149, dev200561 (2022).

47. Wu, J. et al. An alternative pluripotent state confers interspecies chimaeric competency. Nature 521, 316– 321 (2015).

48. Sumi, T., Oki, S., Kitajima, K. & Meno, C. Epiblast ground state is controlled by canonical Wnt/β-catenin signaling in the postimplantation mouse embryo and epiblast stem cells. PLoS One 8, e63378 (2013).

49. Kurek, D. et al. Endogenous WNT signals mediate BMP-induced and spontaneous differentiation of epiblast stem cells and human embryonic stem cells. Stem Cell Reports 4, 114–128 (2015).

50. Edri, S., Hayward, P., Baillie-Johnson, P., Steventon, B. J. & Martinez Arias, A. An epiblast stem cell-derived multipotent progenitor population for axial extension. Development 146, (2019).

51. Najm, F. J. et al. Rapid and robust generation of functional oligodendrocyte progenitor cells from epiblast stem cells. Nat Methods 8, 957–962 (2011).

52. Czechanski, A. et al. Derivation and characterization of mouse embryonic stem cells from permissive and nonpermissive strains. Nature Protocols 9, 559–574 (2014).

53. Mott, R. & Flint, J. Dissecting quantitative traits in mice. Annu Rev Genomics Hum Genet 14, 421–439 (2013).

54. Choi, J. et al. Prolonged Mek1/2 suppression impairs the developmental potential of embryonic stem cells. Nature 548, 219–223 (2017).

55. Mitchell, J. M. et al. Mapping genetic effects on cellular phenotypes with “cell villages”. bioRxiv 2020.06.29.174383 (2020) doi:10.1101/2020.06.29.174383.

56. Neavin, D. R. et al. A village in a dish model system for population-scale hiPSC studies. Nat Commun 14, 3240 (2023).

57. Jerber, J. et al. Population-scale single-cell RNA-seq profiling across dopaminergic neuron differentiation. Nature Genetics 1–9 (2021) doi:10.1038/s41588-021-00801-6.

58. Kang, H. M. et al. Multiplexed droplet single-cell RNA-sequencing using natural genetic variation. Nat Biotechnol 36, 89–94 (2018).

59. Lima, A. et al. Cell competition acts as a purifying selection to eliminate cells with mitochondrial defects during early mouse development. Nat Metab 3, 1091–1108 (2021).

60. Price, C. J. et al. Genetically variant human pluripotent stem cells selectively eliminate wild-type counterparts through YAP-mediated cell competition. Dev Cell 56, 2455–2470.e10 (2021).

61. Macosko, E. Z. et al. Highly Parallel Genome-wide Expression Profiling of Individual Cells Using Nanoliter Droplets. Cell 161, 1202–1214 (2015).

62. Neavin, D. et al. Demuxafy: improvement in droplet assignment by integrating multiple single-cell demultiplexing and doublet detection methods. Genome Biology 25, 94 (2024).

63. Morgan, A. P. et al. The Mouse Universal Genotyping Array: From Substrains to Subspecies. G3 (Bethesda) 6, 263–279 (2015).

64. Broman, K. W. A generic hidden Markov model for multiparent populations. G3 Genes|Genomes|Genetics 12, jkab396 (2022).

65. Lobo, A. K. et al. Identification of sample mix-ups and mixtures in microbiome data in Diversity Outbred mice. G3 (Bethesda) 11, jkab308 (2021).

66. Tabbaa, M., Knoll, A. & Levitt, P. Mouse population genetics phenocopies heterogeneity of human Chd8 haploinsufficiency. 21.

67. Chen, R. et al. Analysis of 589,306 genomes identifies individuals resilient to severe Mendelian childhood diseases. Nat Biotechnol 34, 531–538 (2016).

68. Perrone, F., Cacace, R., van der Zee, J. & Van Broeckhoven, C. Emerging genetic complexity and rare genetic variants in neurodegenerative brain diseases. Genome Medicine 13, 59 (2021).

69. Strano, A., Tuck, E., Stubbs, V. E. & Livesey, F. J. Variable Outcomes in Neural Differentiation of Human PSCs Arise from Intrinsic Differences in Developmental Signaling Pathways. Cell Rep 31, (2020).

70. Schulz, E. G. et al. The Two Active X Chromosomes in Female ESCs Block Exit from the Pluripotent State by Modulating the ESC Signaling Network. Cell Stem Cell 14, 203–216 (2014).

71. Choi, J. et al. DUSP9 Modulates DNA Hypomethylation in Female Mouse Pluripotent Stem Cells. Cell Stem Cell 20, 706–719.e7 (2017).

72. Zvetkova, I. et al. Global hypomethylation of the genome in XX embryonic stem cells. Nat Genet 37, 1274– 1279 (2005).

73. Song, J. et al. X-Chromosome Dosage Modulates Multiple Molecular and Cellular Properties of Mouse Pluripotent Stem Cells Independently of Global DNA Methylation Levels. Stem Cell Reports 12, 333–350 (2019).

74. Tesar, P. J. et al. New cell lines from mouse epiblast share defining features with human embryonic stem cells. Nature 448, 196–199 (2007).

75. Sugimoto, M. et al. A Simple and Robust Method for Establishing Homogeneous Mouse Epiblast Stem Cell Lines by Wnt Inhibition. Stem Cell Reports 4, 744–757 (2015).

76. Takahashi, S., Kobayashi, S. & Hiratani, I. Epigenetic differences between naïve and primed pluripotent stem cells. Cell Mol Life Sci 75, 1191–1203 (2018).

77. Byers, C. et al. Genetic control of the pluripotency epigenome determines differentiation bias in mouse embryonic stem cells. The EMBO Journal 41, e109445 (2022).

78. O’Connor, C. Cell morphology QTL reveal gene by environment interactions in a genetically diverse cell population.

79. Phifer-Rixey, M. & Nachman, M. W. Insights into mammalian biology from the wild house mouse Mus musculus. Elife 4, e05959 (2015).

80. Li, H. & Auwerx, J. Mouse Systems Genetics as a Prelude to Precision Medicine. Trends in Genetics 36, 259–272 (2020).

81. Morgani, S. M., Metzger, J. J., Nichols, J., Siggia, E. D. & Hadjantonakis, A.-K. Micropattern differentiation of mouse pluripotent stem cells recapitulates embryo regionalized cell fate patterning. eLife 7, e32839 (2018).

82. Broman, K. W. et al. R/qtl2: Software for Mapping Quantitative Trait Loci with High-Dimensional Data and Multiparent Populations. Genetics 211, 495–502 (2019).

